# Origin and evolution of the bread wheat D genome

**DOI:** 10.1101/2023.11.29.568958

**Authors:** Emile Cavalet-Giorsa, Andrea González-Muñoz, Naveenkumar Athiyannan, Samuel Holden, Adil Salhi, Catherine Gardener, Jesús Quiroz-Chávez, Samira M. Rustamova, Ahmed F. Elkot, Mehran Patpour, Awais Rasheed, Long Mao, Evans S. Lagudah, Sambasivam K. Periyannan, Amir Sharon, Axel Himmelbach, Jochen C. Reif, Manuela Knauft, Martin Mascher, Nils Stein, Noam Chayut, Sreya Ghosh, Dragan Perovic, Alexander Putra, Ana B. Perera, Chia-Yi Hu, Guotai Yu, Hanin Ibrahim Ahmed, Konstanze D. Laquai, Luis F. Rivera, Renjie Chen, Yajun Wang, Xin Gao, Sanzhen Liu, W. John Raupp, Eric L. Olson, Jong-Yeol Lee, Parveen Chhuneja, Satinder Kaur, Peng Zhang, Robert F. Park, Yi Ding, Deng-Cai Liu, Wanlong Li, Firuza Y. Nasyrova, Jan Dvorak, Mehrdad Abbasi, Meng Li, Naveen Kumar, Wilku B. Meyer, Willem H. P. Boshoff, Brian J. Steffenson, Oadi Matny, Parva K. Sharma, Vijay K. Tiwari, Surbhi Grewal, Curtis Pozniak, Harmeet Singh Chawla, Jennifer Ens, Luke T. Dunning, James A. Kolmer, Gerard R. Lazo, Steven Xu, Yongqiang Gu, Xianyang Xu, Cristobal Uauy, Michael Abrouk, Salim Bougouffa, Gurcharn S. Brar, Brande B. H. Wulff, Simon G. Krattinger

**Affiliations:** Plant Science Program, Biological and Environmental Science and Engineering Division (BESE), King Abdullah University of Science and Technology (KAUST), Thuwal, 23955-6900, Saudi Arabia; Faculty of Land and Food Systems, The University of British Columbia (UBC), Vancouver V6T 1Z4, BC, Canada; Computational Bioscience Research Center (CBRC), King Abdullah University of Science and Technology (KAUST), Computer, Electrical and Mathematical Sciences and Engineering Division (CEMSE), Thuwal, 23955-6900, Saudi Arabia; John Innes Centre, Norwich Research Park, Norwich NR4 7UH, UK; Institute of Molecular Biology and Biotechnology, Azerbaijan National Academy of Sciences, 2a Matbuat Avenue, Baku, 1073, Azerbaijan; Wheat Research Department, Field Crops Research Institute, Agricultural Research Center, 12619, Giza, Egypt; Department of Agroecology, Aarhus University, Slagelse, Denmark; Department of Plant Sciences, Quaid-i-Azam University, Islamabad 45320, Pakistan; International Maize and Wheat Improvement Centre (CIMMYT), c/o CAAS, 12 Zhongguancun South Street, Beijing 100081, China; National Key Facility for Crop Gene Resources and Genetic Improvement & State Key Laboratory of Crop Gene Resources and Breeding, Institute of Crop Science, Chinese Academy of Agricultural Sciences, Beijing 100081, China; Commonwealth Scientific and Industrial Research Organization (CSIRO), Agriculture and Food, Canberra, NSW, Australia; Institute for Cereal Crops Improvement, School of Plant Sciences and Food Security, Tel Aviv University, Tel Aviv, Israel; Leibniz Institute of Plant Genetics and Crop Plant Research (IPK), Gatersleben, Seeland, Germany; Federal Research Centre for Cultivated Plants, Julius Kuehn Institute, Institute for Resistance Research and Stress Tolerance, Quedlinburg, Germany; Bioscience Core Lab, King Abdullah University of Science and Technology (KAUST), Thuwal, 23955-6900, Saudi Arabia; Department of Plant Pathology, Kansas State University, Manhattan, KS, 66506-5502, USA; Department of Plant Pathology and Wheat Genetics Resource Center, Kansas State University, Manhattan, KS, USA; Department of Plant, Soil and Microbial Sciences, Michigan State University, 1066 Bogue Street, Room A286, East Lansing, MI, 48824, USA; National Academy of Agricultural Science, Rural Development Administration, Jeonju, South Korea; School of Agricultural Biotechnology, Punjab Agricultural University, Ludhiana, India; Plant Breeding Institute, University of Sydney, Cobbitty, New South Wales, 2570, Australia; Triticeae Research Institute, Sichuan Agricultural University, Chengdu, Sichuan 611130, China; Department of Biology and Microbiology, South Dakota State University, Brookings, SD, 57007, USA; Institute of Botany, Plant Physiology and Genetics, Tajik National Academy of Sciences, Dushanbe, Tajikistan; Department of Plant Sciences, University of California, Davis, CA, USA; Department of Plant Sciences, University of the Free State, Bloemfontein, South Africa; Department of Plant Pathology, University of Minnesota, Saint Paul, MN, USA; Department of Plant Science and Landscape Architecture, University of Maryland, College Park, MD 20724, USA; Nottingham BBSRC Wheat Research Centre, School of Biosciences, Univ. of Nottingham, Loughborough, UK; University of Saskatchewan, Crop Development Centre, Agriculture Building, 51 Campus Drive, Saskatoon, SK S7N 5A8, Canada; Ecology and Evolutionary Biology, School of Biosciences, University of Sheffield, Western Bank, Sheffield S10 2TN, UK; USDA-ARS, Cereal Disease Laboratory, St. Paul, MN, 55106, USA; USDA-ARS, Crop Improvement and Genetics Research Unit, Western Regional Research Center, 800 Buchanan Street, Albany, CA, 94710, USA; USDA-ARS, Peanut and Small Grains Research Unit, Stillwater, Oklahoma 74075, USA; Present address: The University of Southern Queensland, School of Agriculture and Environmental Science, Centre for Crop Health, Toowoomba, QLD 4350, Australia; Present address: Centre d’anthropobiologie et de génomique de Toulouse (CAGT), Laboratoire d’Anthropobiologie et d’Imagerie de Synthèse, CNRS UMR 5288, Faculté de Médecine de Purpan, 37 Allées Jules Guesde, Bâtiment A, 31000, Toulouse, France; Present address: Department of Plant Science, University of Manitoba, Winnipeg, Manitoba, R3T 2N2, Canada

## Abstract

Bread wheat (*Triticum aestivum*) is a globally dominant crop and major source of calories and proteins for the human diet. Compared to its wild ancestors, modern bread wheat shows lower genetic diversity caused by polyploidisation, domestication, and breeding bottlenecks^1,2^. Wild wheat relatives represent genetic reservoirs, harbouring diversity and beneficial alleles that have not been incorporated into bread wheat. Here, we establish and analyse pangenome resources for Tausch’s goatgrass, *Aegilops tauschii*, the donor of the bread wheat D genome. This new pangenome facilitated the cloning of a disease resistance gene and haplotype analysis across a complex disease resistance locus, allowing us to discern alleles from paralogous gene copies. We also reveal the complex genetic composition and history of the bread wheat D genome, involving previously unreported contributions from genetically and geographically discrete *Ae. tauschii* subpopulations. Together, our results reveal the complex history of the bread wheat D genome and demonstrate the potential of wild relatives in crop improvement.

## Main

Bread wheat (*Triticum aestivum*) is one of the most widely cultivated and most successful crop species worldwide, playing a pivotal role in the global food system. Modern bread wheat shows a remarkable geographical distribution and adaptability to various climatic conditions^1^. Current yield gains, however, might be insufficient to meet future bread wheat demands^3^, calling for concerted efforts to diversify and intensify wheat breeding to further raise yields. Bread wheat is an allohexaploid species (2*n* = 6*x* = 42, AABBDD genome) whose evolution involved the hybridisation of three wild grass species. An initial hybridisation between the A genome donor *T. urartu* (2*n* = 2*x* = 14) and an unknown B genome donor related to the goatgrass *Aegilops speltoides* gave rise to tetraploid wild emmer wheat (*T. turgidum* subsp. *dicoccoides*, 2*n* = 4*x* = 28, AABB genome) 0.5-0.8 million years ago^4^. The second hybridisation event happened between a domesticated tetraploid wheat and the D genome progenitor Tausch’s goatgrass (*Ae. tauschii*; 2*n* = 2*x* = 14, DD genome). This hybridisation that gave rise to bread wheat most likely occurred along the southern shores of the Caspian Sea 8,000-11,000 years ago^5,6^. Polyploidisation and domestication events, like the origin of bread wheat, represent extreme genetic bottlenecks^1,2,7,8^. In the case of bread wheat, recurrent hybridisations with wild wheat relatives and other domesticated wheat species have significantly increased genetic diversity following domestication^2,9-13^. The underlying gene flow contributed to the adaptability of bread wheat to diverse climatic conditions outside the Fertile Crescent, the geographical region where wheat was domesticated. In comparison to the A and B genomes, however, D genome diversity in bread wheat remains low because the above gene flow has predominantly involved tetraploid species with an AB genome^2,6,10,14^.

Here, we establish a comprehensive set of pangenome resources for the bread wheat D genome progenitor *Ae. tauschii,* including chromosome-scale assemblies representing the three *Ae. tauschii* lineages and whole-genome sequencing data of a large *Ae. tauschii* diversity panel. The genomic resources proved useful for haplotype and gene discovery and allowed us to unravel the composition and evolution of the bread wheat D genome.

## Results

### The *Aegilops tauschii* pangenome

To comprehensively assess genetic diversity in *Ae. tauschii*, we compiled a presence/absence *k*-mer matrix from a diversity panel comprising 920 sequenced *Ae. tauschii* accessions (Supplementary Table 1, Supplementary Note 1). We optimized the *k*-mer matrix workflow for large diversity panels (Supplementary Note 2) using whole-genome sequencing data from this and previous studies^14,15^ (Supplementary Table 1). The *k*-mer analysis revealed 493 non-redundant (NR) *Ae. tauschii* accessions, whereas the remaining accessions shared at least 96% of their *k*-mers with a given NR accession (Supplementary Table 2). The NR diversity panel spanned the geographical range of *Ae. tauschii* from north-western Turkey to eastern China (Fig. 1a) and defined a phylogeny demarcated by the three basal lineages, with 335 accessions for Lineage 1 (L1), 150 accessions for Lineage 2 (L2), and eight accessions for Lineage 3 (L3) (Fig. 1b). Based on phylogeny (Fig. 1b, Extended Data Fig. 1a) and STRUCTURE analyses (Extended Data Fig. 2), we defined four geographically distinct subpopulations for *Ae. tauschii* L2, referred to as L2E-1 (south-western Caspian Sea), L2E-2 (south-eastern Caspian Sea), L2W-1 (Caucasus), and L2W-2 (Turkmenistan and northern Iran), in accordance with the literature^5,15^. Group L2E from the southern Caspian Sea (representing subpopulations L2E-1 and L2E-2 here) was previously identified as the main contributor of the bread wheat D genome^5^. Of the 150 NR *Ae. tauschii* L2 accessions, we could assign 133 to one of the four L2 subpopulations based on an ancestry threshold ≥70% (Supplementary Table 3). The remaining 17 L2 accessions were considered admixed.

**Fig. 1.**
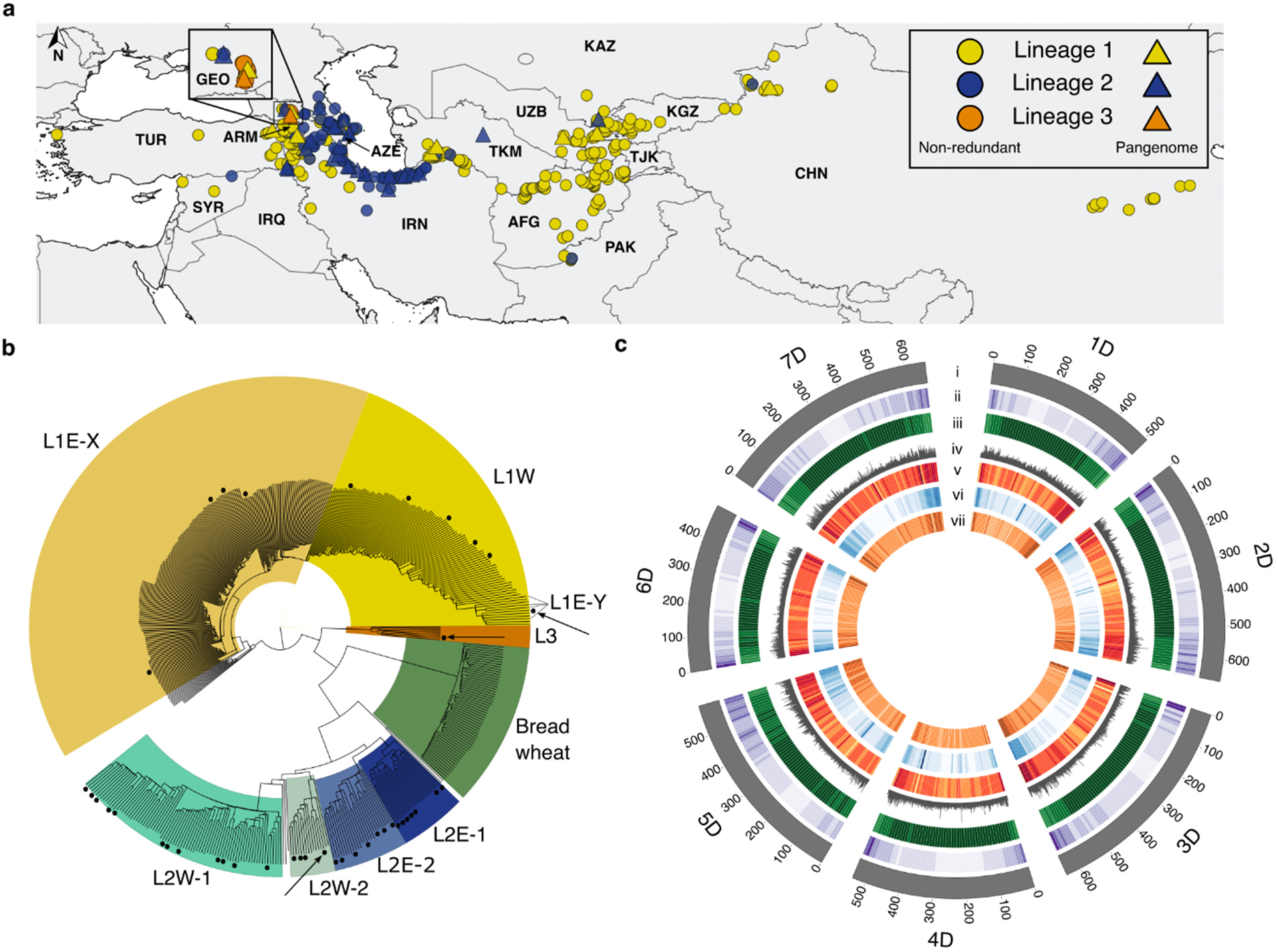
The *Aegilops tauschii* diversity panel and pangenome. **a**, Geographical distribution of the 493 non-redundant *Ae. tauschii* accessions in the diversity panel. Lineage 1 (L1) accessions are indicated in yellow, Lineage 2 (L2) accessions in blue, and Lineage 3 (L3) accessions in orange. The 46 accessions representing the pangenome are indicated by triangles coloured according to lineage. **b**, Phylogeny of the non-redundant *Ae. tauschii* accessions showing the subpopulations within the three lineages as labelled on the tree. Accessions representing the pangenome are indicated by black dots next to the tree branches. The three reference accessions TA10171 (L1), TA1675 (L2) and TA2576 (L3) are indicated by black arrows. The D subgenome from 59 wheat landraces is shown in relation to *Ae. tauschii*. **c**, Circos plot showing annotation features, nucleotide diversity, and structural variants across the *Ae. tauschii* panel and pangenome relative to the TA1675 L2 reference assembly. The tracks show (i) the chromosomes and their length, (ii) gene density for TA1675 (11 – 326 high-confidence genes per 10 Mb window), (iii) repeat density for TA1675 (1,047,826 – 9,723,453 repeat-masked bases per 10 Mb window), (iv) nucleotide diversity across the panel of 493 non-redundant accessions (π = 10*10^-09^ – 0.005 per 10 kb window), and structural variants (SV) density across 10 kb windows in (v) L1 (0 – 9,237 SVs), (vi) L2 (0 – 24,585 SVs) and (vii) L3 (0 – 767 SVs) accessions of the pangenome.

Using genetic, geographical and phenotypic diversity, we selected 46 accessions for pangenome analysis, comprising eleven L1 accessions, 34 L2 accessions and one L3 accession (Fig. 1a, b, Supplementary Table 4). The 46 pangenome accessions captured 99.3% of the genetic diversity present in the *Ae. tauschii* diversity panel based on *k*-mer analysis (Extended Data Fig. 1b, Supplementary Table 5). We sequenced the selected 46 pangenome accessions by PacBio circular consensus sequencing^16^ to a median genome coverage of 23-fold (18-47-fold) and generated primary contig-level assemblies with contig N50 values ranging from 15.02 Mb to 263.79 Mb (median 45.26 Mb) (Supplementary Table 6). Phred quality scores ranged from 34.9 to 48.3 (median 45.5) and *k*-mer completeness scores from 95.1-99.8% (median 99.4%) based on 21-mer content comparison with short-read WGS data (Supplementary Table 6). We calculated the benchmarking universal single-copy orthologs (BUSCO) scores for each accession^17^, returning values between 98.0% and 98.6% (Supplementary Table 6), indicating high contiguity, accuracy, and completeness of the assemblies. We selected one representative accession per lineage to generate *de novo* annotated pseudochromosome assemblies, namely TA10171 (L1), TA1675 (L2), and TA2576 (L3). For these three accessions, we increased the sequencing coverage to 67-97-fold, generated assemblies with contig N50 values of 53.38 Mb (L1), 221.04 Mb (L2), and 116.91 Mb (L3) (Supplementary Table 6), and used Hi-C chromatin conformation capture^18^ to scaffold the assemblies into pseudomolecules (Extended Data Table 1, Extended Data Fig. 3a-c). We then scaffolded the remaining 43 L1 and L2 contig-level assemblies using their respective L1 and L2 chromosome-scale references as guides (Supplementary Table 6).

In accordance with previous observations^5,14,19^, we detected increased nucleotide diversity in L2 (π = 0.00038) compared to L1 and L3 (L1, π = 0.00021; L3, π = 0.00024) (Fig. 1c; Supplementary Table 7). Structural variants (SVs) were called across the pangenome relative to the TA1675 (L2) reference assembly (Fig. 1c). L1 accessions showed a similar distribution of SVs to that in the L3 accession TA2576 (Extended Data Fig. 1c), with a median of 205,856 SVs and 191,179 SVs per accession for L1 and L3, respectively (Supplementary Table 8). The L2 accessions had a median of 85,401 SVs per accession compared to the TA1675 reference (Supplementary Table 8).

### The *Ae. tauschii* pangenome facilitates gene discovery

The highly contiguous *Ae. tauschii* pangenome assemblies generated here present an opportunity for gene discovery and characterisation by comparative haplotype analyses. Here, we assessed the value of the *Ae. tauschii* genomic resources with a focus on rust resistance genes. The three fungal wheat rust diseases, leaf rust (caused by *Puccinia triticina*, *Pt*), stripe rust (*P. striiformis* f. sp. *tritici, Pst*), and stem rust (*P. graminis* f. sp. *tritici*, *Pgt*), are among the most devastating and most ubiquitous wheat diseases, causing considerable yield losses^20^. The stem rust resistance gene *SrTA1662* was introgressed into bread wheat from *Ae. tauschii* accession TA1662 and genetically mapped to the stem rust resistance locus *SR33* on chromosome arm 1DS^21,22^. Because the original mapping could not establish whether the stem rust resistance gene from TA1662 was a new gene or was allelic to *Sr33*, the gene was given the temporary designation *SrTA1662*^21^. *Sr33* and *SrTA1662* encode intracellular nucleotide-binding leucine-rich repeat (NLR) immune receptors belonging to the *Mla* family (Fig. 2b, Extended Data Fig. 4a)^14,22,23^. Here, we repeated the *k*-mer-based association mapping that led to the initial discovery of the *SrTA1662* candidate gene^14^. Compared to the short-read based *Ae. tauschii* assemblies^14^, mapping the *k*-mers against our high-quality *Ae. tauschii* genome strongly decreased the noise in the *k*-mer-based association approach (Fig. 2a). A detailed haplotype analysis revealed that *SrTA1662* is a paralogue rather than an allele of *Sr33* (Fig. 2b, Extended Data Fig. 4a, Supplementary Table 9). When we compared the stem rust infection phenotypes of *Ae. tauschii* lines predicted to carry only *Sr33* or *SrTA1662*, we observed that the two genes appeared to have different specificities (Supplementary Table 10). We confirmed this notion by inoculating *SrTA1662* transgenic wheat lines^14^ and *Sr33* introgression lines with three *Pgt* isolates (Extended Data Fig. 4b, Supplementary Table 11). In line with the nomenclature standards for wheat gene designation^24^, we therefore renamed *SrTA1662* as *Sr66*.

**Figure 2.**
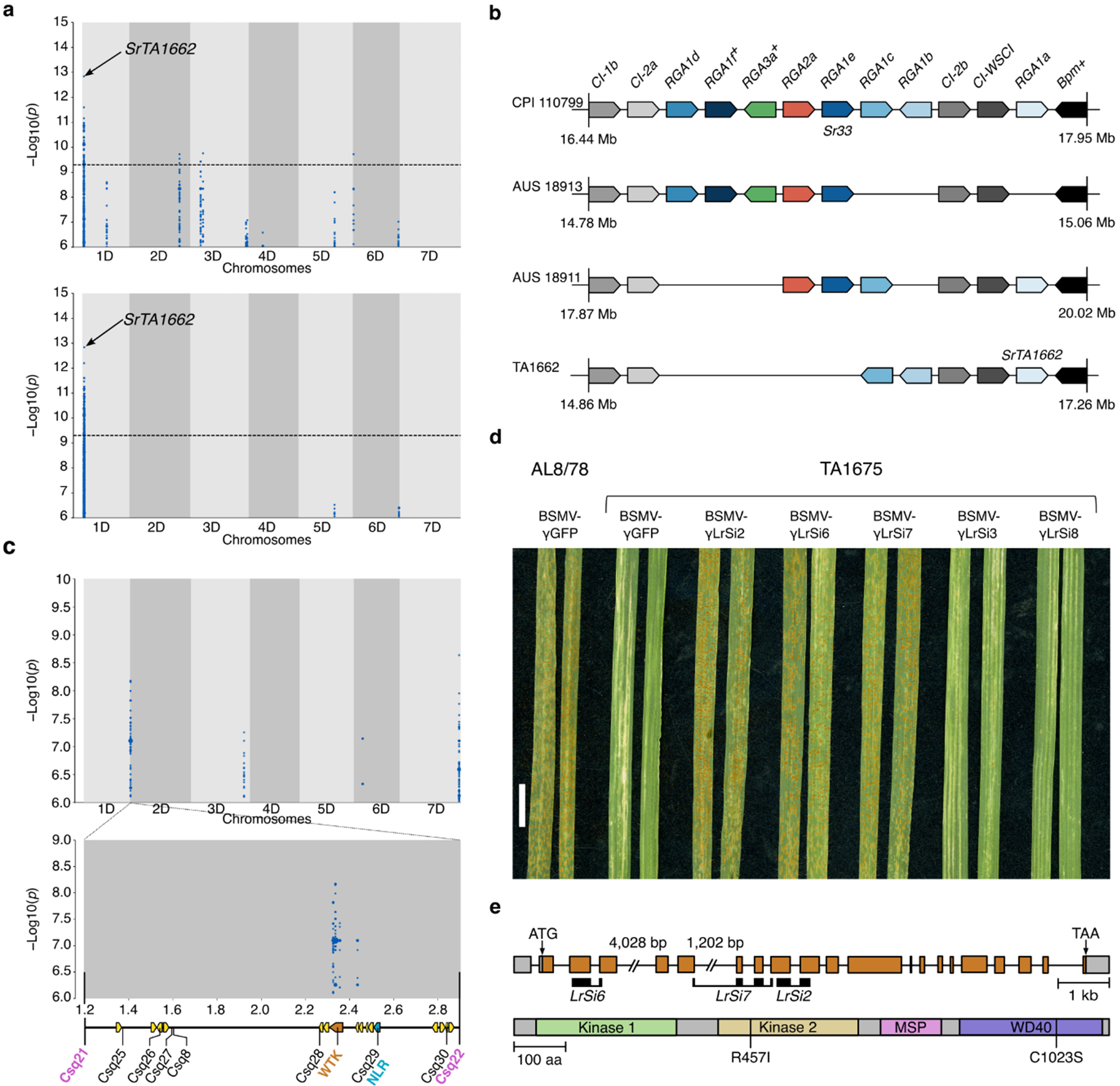
Haplotype analyses and cloning of a disease resistance gene using the *Ae. tauschii* pangenome. **a**, Effect of assembly quality on association genetics. Significantly associated *k*-mers for resistance to *Puccinia graminis* f. sp. *tritici* (*Pgt*) race QTHJC mapped to *Ae. tauschii* accession TA1662 using a low-quality assembly (top, contig N50 of 196 kb)^14^ with contigs anchored to the chromosome-scale TA1675 reference assembly, and a high-quality TA1662 assembly (bottom, contig N50 of 58.21 Mb, this study) scaffolded against the TA1675 reference assembly. The significantly associated peak in chromosome arm 1DS corresponds to the linkage disequilibrium (LD) block containing the stem rust resistance gene *SrTA1662* (*Sr66*). **b**, Haplotype analysis across the *Mla* locus of chromosome arm 1DS. Shown are Resistance Gene Analogues (*RGA*) in *Ae. tauschii* accessions CPI 110799, AUS 18913, AUS 18911 and TA1662. Boxes indicate genes and their directionality indicates the direction of transcription. Pseudogenes are indicated by a superscripted plus sign after the gene name. *RGA* alleles across accessions are indicated by the same colour and position. The *RGA* locus is flanked by subtilisin-chymotrypsin inhibitor (*CI*) genes (grey) and a pumilio (*Bpm*) homologous gene (black). Other unrelated genes present in this region are omitted. The locus length and distribution of the genes are not drawn to scale. **c**, *k*-mer-based genome-wide association with an isolate of *P. triticina* race BBBDB mapped to the chromosome-scale assembly of *Ae. tauschii* accession TA1675. The peak on the short arm of chromosome 2D corresponds to the leaf rust resistance locus *LR39*. The diagram shows the *LR39* target interval in *Ae. tauschii* accession TA1675. *Csq21* and *Csq22* represent markers flanking *LR39* identified through bi-parental mapping. Markers *Csq8* and *Csq25*-*Csq30* co-segregated with *LR39*. Arrows indicate candidate genes. *WTK*, wheat tandem kinase gene; *NLR*, nucleotide-binding leucine-rich repeat gene. **d**, Representative images showing the results of virus-induced gene silencing (VIGS). AL8/78, susceptible control; BSMV-γGFP, barley stripe mosaic virus (BSMV) expressing a GFP silencing construct (control); BSMV-γLrSi2, BSMV-γLrSi6, and BSMV-γLrSi7 are silencing constructs specific for the *WTK* gene. BSMV-γLrSi3 and BSMV-γLrSi8 are silencing construct specific for the *NLR* gene. The specificities of the silencing constructs were evaluated using the TA1675 chromosome-scale assembly. Chlorotic areas on the BSMV-γGFP controls represent virus symptoms. Scale bar = 1 cm. **e**, Gene structure of *Lr39* and domain architecture of the Lr39 protein. Grey boxes, untranslated regions; orange boxes, exons; lines, introns. The positions of the VIGS silencing probes are indicated. R457I and C1023S in the protein structure indicate the two Lr39 amino acid changes between TA1675 (resistant) and AL8/78 (susceptible).

*Ae. tauschii* accession TA1675, for which a chromosome-scale reference assembly was generated in this study, is the donor of the leaf rust resistance gene *Lr39*, which was mapped to the short arm of chromosome 2D^25,26^. *k*-mer-based association mapping with the *Pt* isolate BBBDB (avirulent against *Lr39*)^27^ revealed a peak at the telomeric end of chromosome arm 2DS, corresponding to the 2.33-2.45 Mb region in the TA1675 assembly (Fig. 2c). This location overlapped with markers flanking *LR39* (positions 1.20-2.84 Mb) that were identified based on bi-parental genetic mapping (Fig. 2c, Extended Data Fig. 5a, b)^28^. The genomic region underlying the association peak contained three candidate genes in TA1675 (one wheat tandem kinase and two genes of unknown function), while the interval identified through bi-parental mapping harboured 16 genes (Supplementary Table 12). Based on functional annotations and polymorphisms compared to the susceptible *Ae. tauschii* accession AL8/78, the most promising candidate was *AeT.TA1675.r1.2D000150*, encoding a wheat tandem kinase (WTK), a protein family that plays a prominent role in disease resistance in wheat^29-31^. Virus-induced gene silencing (VIGS) of the *WTK* candidate gene in TA1675 resulted in greater susceptibility to leaf rust (Fig. 2d). Silencing of an NLR-encoding gene (*AeT.TA1675.r1.2D000200*) located just outside the peak region did not result in increased susceptibility (Fig. 2 c, d), indicating that the *WTK* gene is *Lr39*. The predicted genomic sequence of the *Lr39* candidate gene is 11,699 bp in length with 21 exons. The corresponding 3,408-bp coding sequence encodes an 1,135-amino acid protein with two N-terminal kinase domains of the LRR_8B subfamily, followed by a major sperm protein (MSP) domain and a WD40 repeat-containing domain (WD40) at the C terminus (Fig. 2e, Extended Data Fig. 5c). Compared to the susceptible *Ae. tauschii* accession AL8/78, Lr39 from TA1675 carried two amino acid changes located in the kinase 2 and WD40 domains, respectively (Fig. 2e, Extended Data Fig. 5c).

### Tracing lineage-specific *Ae. tauschii* haplotype blocks in the bread wheat genome

Bread wheat has become one of the most successful and widely cultivated crop species that is adapted to a wide range of climatic conditions^1^. Continuous gene flow by natural and artificial introgressions from wild wheat relatives has increased the genetic diversity of bread wheat following a domestication bottleneck^9-13^. For example, around 1% of the extant bread wheat D genome originated from *Ae. tauschii* L3, indicating multiple hybridisation events that gave rise to the extant bread wheat D genome^14^. We hypothesise that bread wheat landraces with higher L3 genome content, possibly representing a more ancestral state of the L3 introgression(s), have been preserved in *ex situ* collections but are rare and geographically restricted.

To identify bread wheat accessions with above-average proportions of L3 genome, we developed the ‘Missing Link Finder’ pipeline (Fig. 3a). Missing Link Finder estimates the similarity between a species- or lineage-specific reference *k*-mer set and sample *k*-mer sets generated from genotyping data of individual wheat accessions, computing the result as Jaccard similarity coefficients. To deploy Missing Link Finder, we used a reference *k*-mer set consisting of 769 million *Ae. tauschii* L3-specific *k*-mers^14^ and compared it to individual sample *k*-mer sets from 82,293 genotyped wheat accessions (6.16 million *k*-mers per accession on average)^32^. We identified 503 bread wheat accessions with an above-average (>2-fold) normalized Jaccard index (a value of 0 indicates average number of L3 *k*-mers), indicative of increased *Ae. tauschii* L3 content (Fig. 3b, Extended Data Fig. 6a). The 139 accessions with the highest Jaccard indices are synthetic hexaploid wheats (Extended Data Fig. 6a), most of which (122 accessions) were produced using an *Ae. tauschii* accession collected in Georgia (CWI 94855), the only country where *Ae. tauschii* L3 has been found in the present day^14^. We also identified 364 bread wheat landraces with putatively increased proportions of L3 introgressions (Fig. 3b, Extended Data Fig. 6a). One of the bread wheat landraces with the highest Jaccard indices, CWI 86942 (PI 572674), was collected in the Samegrelo-Zemo Svaneti region of Georgia^33^. We observed a gradient of decreasing L3 proportions (as revealed by Jaccard indices) with increasing geographical distance from Georgia (Fig. 3c, Extended Data Fig. 6b).

**Fig. 3.**
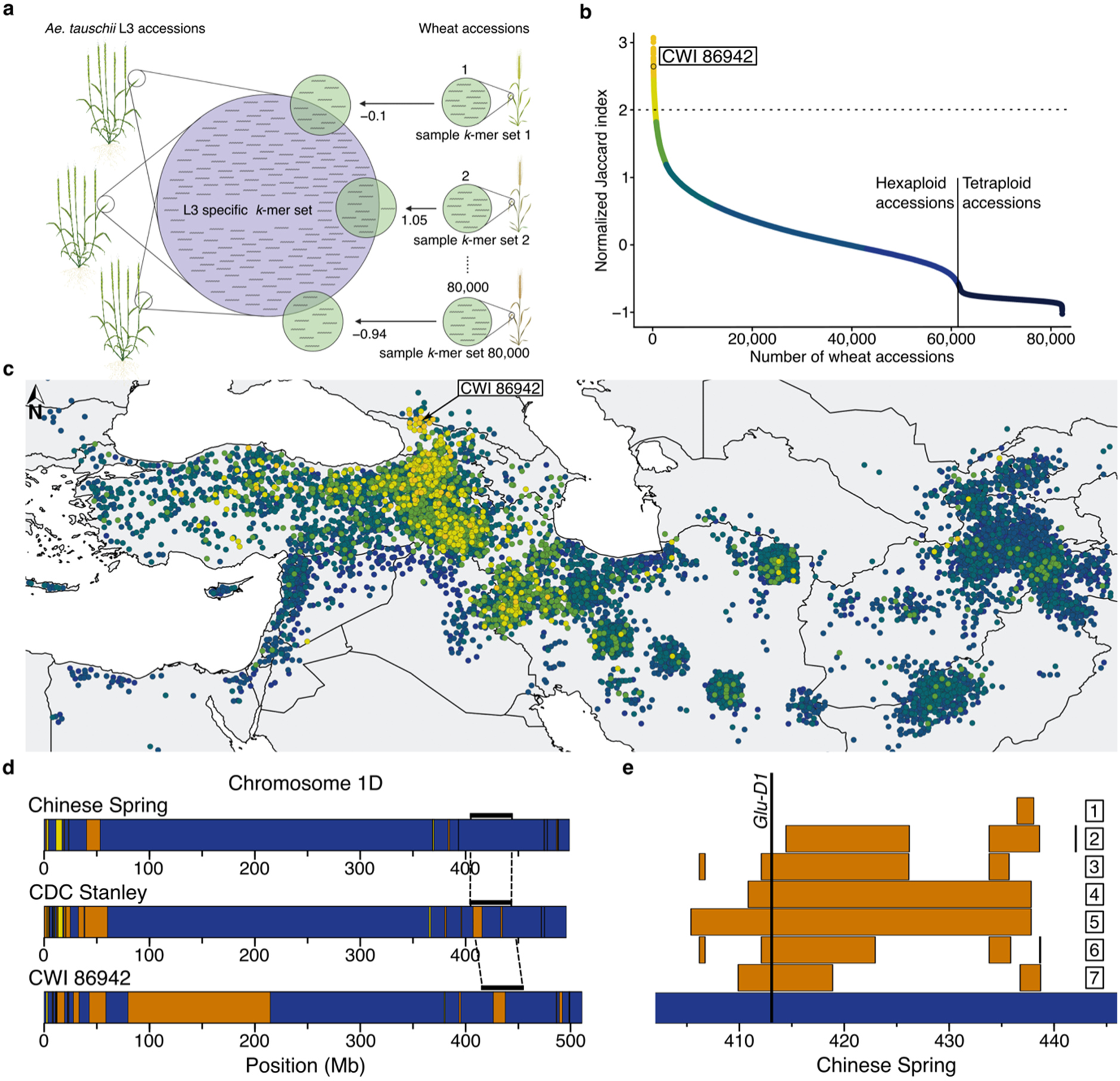
*Ae. tauschii* L3 introgressions in bread wheat. **a**, Diagram of the Missing-Link Finder pipeline. An *Ae. tauschii* L3-specific *k*-mer set (769 million L3-specific *k*-mers; blue circle) was compared to individual sample *k*-mer sets generated from over 80,000 genotyped wheat accessions (green circles). The result is indicated as normalized Jaccard indices. **b**, Distribution of normalized Jaccard scores across 82,154 wheat accessions. The horizontal dotted line indicates the >2 threshold. The 139 synthetic hexaploid wheat lines with increased Jaccard indices have been removed and are shown in Extended Data Fig. 6a. **c**, The Jaccard indices show a gradual decline with increasing geographical distance from Georgia. Dots represent individual bread wheat accessions for which exact coordinates were available. Colours represent different normalized Jaccard indices corresponding to Fig. 3b. A full map is shown in Extended Data Fig. 6b **d**, Diagram of chromosome 1D in the wheat lines Chinese Spring, CDC Stanley, and CWI 86942. Haplotype blocks corresponding to *Ae. tauschii* L1 are indicated in yellow, L2 in blue, and L3 in orange. The black bars above the chromosome indicate the region shown in (**e**). **e**, Diagram of a portion of the long arm of bread wheat chromosome 1D. Shown are different lengths of the L3 introgression segment in various bread wheat lines. The numbers correspond to the following accessions chosen for their diverse recombination patterns in this locus: 1, CWI 86929; 2, CWI 30140; 3, CWI 57175; 4, CWI 84686; 5, CWI 84704; 6, CWI 86481; 7, CDC Stanley.

To further quantify and explore the L3 contents of CWI 86942 and other landraces, we generated an annotated chromosome-scale assembly of CWI 86942 using PacBio circular consensus sequencing^16^ and chromosome conformation capture^18^ (Extended Data Table 1, Extended Data Fig. 3d). In addition, we produced whole-genome sequencing data (10-fold coverage) of 36 hexaploid wheat landraces with higher (>2) Jaccard indices using short-read Illumina-based sequencing. For comparison, we also sequenced 23 wheat landraces with Jaccard indices <2 (Supplementary Table 13). Our analysis focused on landraces to avoid detection of L3 haplotype blocks that might be the result of artificial introgressions. We observed a good correlation between the Jaccard indices and the *Ae. tauschii* L3 content estimated based on whole-genome sequencing data (Extended Data Fig. 6c), supporting the idea that Missing Link Finder is a suitable pipeline to identify rare wheat accessions with above-average introgressions. CWI 86942 contained ∼7.0% of L3 introgressions, compared to the 0.5%-1.9% in other bread wheat assemblies^14^. Most notable was a 135-Mb L3 segment in the pericentromeric region of chromosome 1D (Fig. 3d), which represents the largest *Ae. tauschii* L3 haplotype block reported in bread wheat so far. This segment contains 587 predicted genes, of which 112 showed presence-absence variation or a disruptive mutation compared to the corresponding L2 segment in wheat cultivar Kariega (Supplementary Table 14). In addition to CWI 86942, this L3 haplotype segment, or parts thereof, were found in multiple bread wheat landraces collected between the 1920s and the 1930s (Extended Data Fig. 6d), indicating that this segment is not the result of synthetic hexaploid wheat breeding^34,35^. A second notable L3 segment was found on the long arm of chromosome 1D in multiple bread wheat landraces (Fig. 3e). This segment carries a superior wheat quality allele at the *Glu-D1* locus that originated from *Ae. tauschii* L3^36^. In modern bread wheat (for example, wheat cultivars CDC Landmark, CDC Stanley, and Jagger), this L3 segment is ∼8.5 Mb in size. We identified a group of bread wheat landraces originating from Azerbaijan where the corresponding L3 segment was up to 36.35 Mb in size (Fig. 3e). This L3 introgression showed various lengths in different bread wheat landraces, reflecting extensive recombination. We further estimated the cumulative proportion of L3 introgression across a comprehensive set of 126 hexaploid wheat landraces, including the whole-genome sequencing data from the 59 landraces generated in this study and publicly available sequencing data (Supplementary Table 15). Using identity-by-state, we determined that 16.6% of the wheat D genome, corresponding to 666.0 Mb and containing 8,779 high-confidence genes (25.6%), can be covered with *Ae. tauschii* L3 haplotype blocks across these landraces (Supplementary Table 16). Although the proportion of *Ae. tauschii* L3 introgressions in most modern bread wheat cultivars is marginal (∼1% relative to the entire D genome), the cumulative size of L3 introgressions across multiple bread wheat landraces is considerable.

### Origin and evolution of the wheat D genome

We determined the complexity and origin of the D genome across 17 hexaploid wheat lines, for which chromosome-scale assemblies are available^11,37-42^. We divided the wheat genomes into 50-kb windows and assigned each window to an *Ae. tauschii* subpopulation based on identity-by-state^13^. We observed that all four *Ae. tauschii* L2 subpopulations contributed genomic segments to the bread wheat D genome (Supplementary Table 17). Consistent with previous reports^5^, the largest proportion of the wheat D genome (45.6-51.3%) originated from subpopulation L2E-1, which is mainly found in the south-western Caspian Sea region. Subpopulation L2E-2 (south-eastern Caspian Sea) contributed 24.7-27.3% to the wheat D genome (Fig. 4a). Up to 6.9% of the wheat D genome was identical (based on identity-by-state analysis) to *Ae. tauschii* subpopulation L2W, with 4.1-5.0% coming from subpopulation L2W-1, and subpopulation L2W-2 contributing 1.7-2.2% (Fig. 4a, b, Extended Data Fig. 7). We could assign another 10.7-19.5% of the wheat D genome to L2, but without being able to infer the exact subpopulation, indicating that these segments originated from *Ae. tauschii* L2 haplotypes that were not captured in our diversity panel (Fig. 4a, b, Extended Data Fig. 7). The contributions from *Ae. tauschii* L1 and L3 ranged from 0.7% to 1.1% and 1.6% to 7.0%, respectively. Genomic windows representing 0.1-2.4% of the hexaploid wheat D genome had a different origin than *Ae. tauschii*. These windows include previously described introgressions, like the tall wheatgrass (*Thinopyrum ponticum*) introgression on chromosome 3D in bread wheat cultivar LongReach Lancer or the putative *Ae. markgrafii/Ae. umbellulata* introgression on chromosome 2D of cultivars Julius, Arina*LrFor*, SY Mattis, and Jagger^11,12^ (Supplementary Table 17). The number of *Ae. tauschii* subpopulations that contributed to the hexaploid wheat D genome does not necessarily reflect the number of independent hybridization events, because the *Ae. tauschii* line that contributed the D genome may have already been admixed. To infer the minimal number of hybridisations that gave rise to the extant wheat D genome, we assessed the number of haplotypes present at any given position in the wheat D genome. We used Chinese Spring as a reference and identified 50-kb windows showing no identity-by-state across the 126 hexaploid wheat landraces for which whole-genome sequencing data were available. Such windows indicate the presence of at least two haplotypes in the hexaploid wheat gene pool. Consecutive 50-kb windows with no identity-by-state were concatenated into alternative haplotype blocks. The origins of alternative haplotype blocks were then assigned to one of the *Ae. tauschii* subpopulations using identity-by-state (Supplementary Table 18). In total, 71.4% of the wheat D genome was covered by a single haplotype across the analysed hexaploid wheat lines (59.7% of genes). The remaining 28.6% of the wheat D genome showed multiple haplotypes (21.0% of the wheat D genome had two haplotypes [26.2% of genes], 6.7% had three haplotypes [12.2% of genes], and 0.9% four haplotypes [1.9% of genes]) (Fig. 4b, Extended Data Fig. 7, Supplementary Table 19). The maximum number of haplotypes corresponding to different *Ae. tauschii* subpopulations for any given window was four, indicating that the bread wheat D genome evolved through at least four hybridisations.

**Fig. 4.**
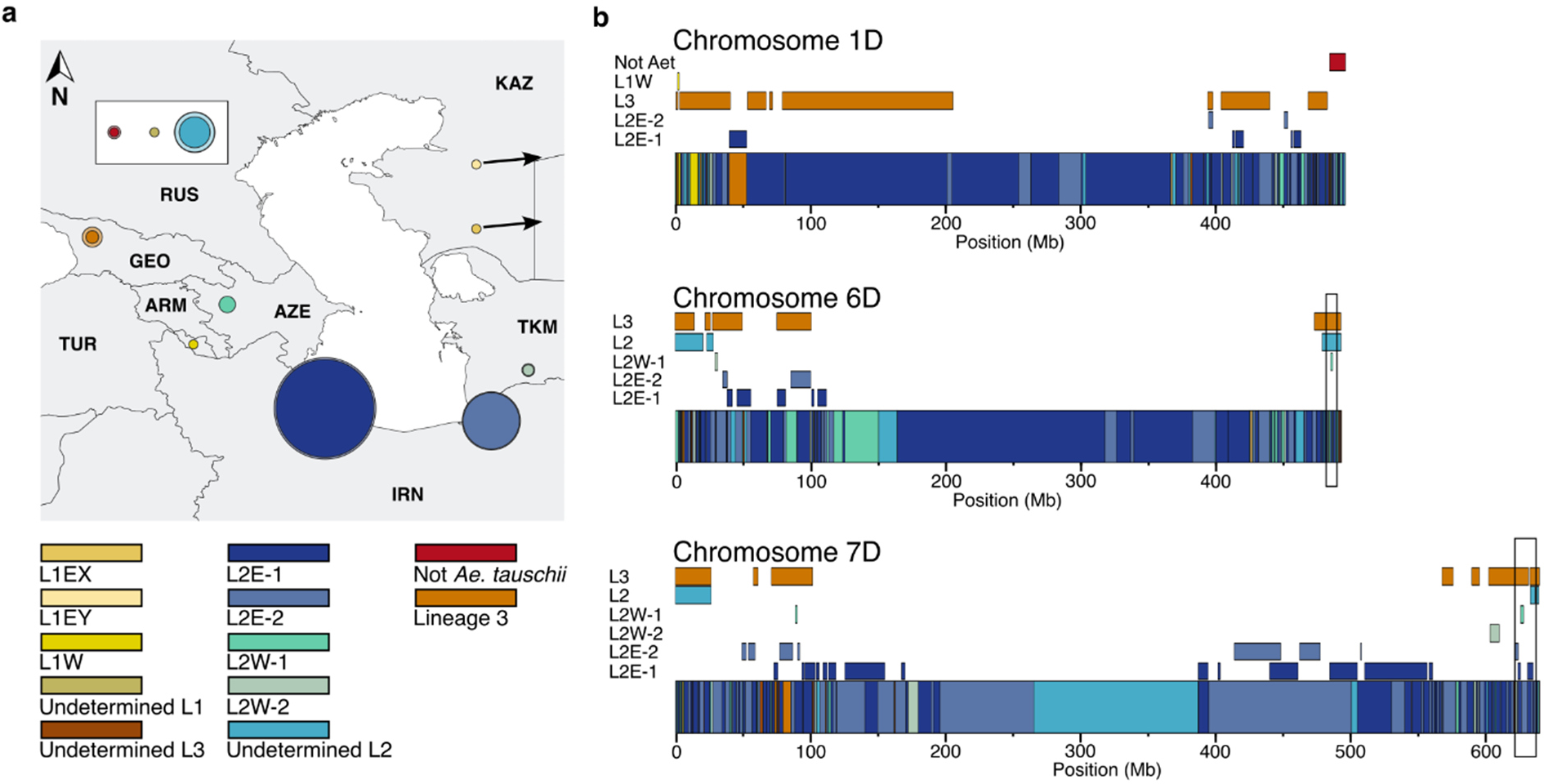
Different *Ae. tauschii* subpopulations contributed to the hexaploid wheat D genome. **a**, Proportions of *Ae. tauschii* subpopulations that make up the wheat D genome. Inner circles in solid colours represent the average proportions across 17 hexaploid wheat assemblies. The outer lighter circles represent the maximum proportion found across the 17 wheat genomes. The geographical location for each subpopulation was assigned based on representative accessions. **b**, Minimal number of hybridisation events that gave rise to the extant bread wheat D genome. Diagrams of chromosomes 1D, 6D, and 7D in Chinese Spring. The coloured boxes along the chromosomes represent the haplotypes present in Chinese Spring. Coloured rectangles above the chromosomes represent alternative haplotype blocks identified across 126 hexaploid wheat landraces (cumulative length of alternative haplotype blocks across all 126 landraces). Colours refer to the *Ae. tauschii* subpopulations. The maximum number of haplotype blocks was four. Black boxes highlight the regions on chromosome 3D and 7D in which four overlapping haplotypes are found.

## Discussion

The comprehensive genomic resources generated in this study enabled haplotype analysis and cloning of rust resistance genes and they offered a detailed insight into the composition and origin of the bread wheat D genome. Crop domestication has often been considered as a relatively simple linear progression^43^. Our analyses support a model of protracted domestication that is more complex, involving recurring episodes of hybridisation and gene flow that resulted in patchwork-like haplotype patterns across the bread wheat D genome. We largely confirm that an *Ae. tauschii* L2 population from the south-western Caspian Sea region was the major donor of the bread wheat D genome (Fig. 4a)^5^, with smaller genomic segments originating from different *Ae. tauschii* lineages^5,14^. In contrast to previous reports, however, our work revealed a much more complex patterning of the bread wheat D genome. We determined that all four L2 subpopulations, as well as L1 and L3, contributed segments to the extant bread wheat D genome. Compared to the AB subgenomes, the bread wheat D genome shows a lower genetic diversity, indicative of a much lower rate of introgression from wild progenitors^2,6^. The patchwork pattern seen in the bread wheat D genome is somewhat surprising given that most *Ae. tauschii* L2 accessions in our diversity panel showed a low degree of admixture, with a well-defined population structure following their geographical distribution (Extended Data Fig. 2). A possible explanation for this observation is that the *Ae. tauschii* accession that gave rise to the bread wheat D genome was admixed, carrying genomic segments from different subpopulations.

Remnants of *Aegilops* species have been identified at several pre-agricultural settlements in the Fertile Crescent^44^, indicating that *Aegilops* species were used as food source or persisted as weeds in pre-agricultural cultivation of other wild cereals. The gathering and possible management of *Ae. tauschii* for food, or its co-cultivation as a weed over an extended period might have resulted in mixing of *Ae. tauschii* populations with different geographical origins, leading to an increase of admixed accessions close to human settlements. Such an admixed *Ae. tauschii* population might have later given rise to the bread wheat D genome. This scenario would also explain why the bread wheat D genome forms a separate clade from *Ae. tauschii* in many phylogenetic and population structure analyses (Fig. 1b)^5,14,15,45^. Alternatively, the *Ae. tauschii* accession that gave rise to the bread wheat D genome was non-admixed, and recurrent hybridisations resulted in the observed mosaic-like haplotype pattern.

Another important finding of this study is the large cumulative size of alternative haplotype (non-L2E) blocks in the bread wheat D genome. Following hexaploidisation, genetic material from the other *Ae. tauschii* lineages (L1 and L3) became incorporated into the bread wheat D genome and were subsequently broken into smaller fragments via recombination. Although the proportions of alternative haplotype blocks are low in individual elite wheat cultivars, the different segments accumulate to considerable lengths across various genotypes. This notion is evidenced by the cumulative size of L3 segments that span a total of 666.0 Mb. We assessed 126 hexaploid wheat landraces, and, although we selected for accessions with increased L3 genome content, it is likely that the proportion of remnant L3 segments in the bread wheat gene pool is even higher. This finding raises important questions about the adaptive potential of alternative haplotype blocks for wheat breeding.

## Acknowledgements

This research used the Shaheen III supercomputer and the Ibex cluster managed by the Supercomputing Core Laboratory at King Abdullah University of Science and Technology (KAUST) in Thuwal, Saudi Arabia. We thank the systems administrators and computational scientists for help with debugging and overall support. We are grateful to Burkhard Steuernagel (JIC), David Keyes, Leonardo Fabbian and Kenyi González Kise (all KAUST) for bioinformatics advice, Elisabet Poquet Faig (KAUST) for greenhouse assistance, Ines Walde (IPK) for technical assistance with Hi-C library preparation and sequencing, Guo (Cherry) Huijuan (Novogene) for administering the OWWC sequencing, Jesse Poland (KAUST) and Liangliang Gao (KSU) for advice on PacBio sequencing, Allison Bentley (CIMMYT), Beat Keller (University of Zurich), Herman Bürstmayr (BOKU), Justin Faris (USDA), Marco Maccaferri and Matteo Bozzoli (both UNIBO), Richard Horsnell (NIAB), David Seung, Janneke Balk, Rose McNelly, and Tom O’Hara (all JIC) for nominating *Ae. tauschii* lines of strategic interest, Bob McIntosh (University of Sydney) for critical reading of the manuscript, and Emma Waller for assistance with the OWWC website. This publication is based upon work supported by KAUST awards ORFS-CRG10-2021-4735 to B.B.H.W., ORFS-CRG11-2022-5076 to S.K., and URF/1/4352-01-01, FCC/1/1976-44-01, FCC/1/1976-45-01, REI/1/5234-01-01, and REI/1/5414-01-01 to X.G.; Australian Government Research Training Program and the University of Queensland Centennial Scholarships to N.A. during genetic mapping of *LR39*; Academy of Scientific Research and Technology Project ID 19385 and Climate Change Adaptation and Nature Conservation (GREEN FUND) to A.F.E.; National Major Agricultural Science and Technology project NK2022060101 and National Key Research and Development Program of China 2021YFF1000204 to L.M.; a Genbank3.0 project from the German Federal Ministry of Education and Research (grant no. FKZ 031B1300A) to J.C.R.; the UK Biotechnology and Biological Sciences Research Council (BBSRC) Institute Strategic Programme Designing Future Wheat (BB/P016855/1) and a European Research Council grant (ERC-2019-COG-866328) to C.U.; the Mexican Consejo Nacional de Ciencia y Tecnología (CONACYT; 2018-000009-01EXTF-00306) to J.Q.-C.; funding from Department of Biotechnology, Government of India to P.C. and S.K.; USDA-NIFA Capacity Fund via South Dakota Agricultural Experiment Station to W.L.; a Bayer (previously Monsanto) Beachell Borlaug International Scholar’s Program to S.G.; National Science Foundation of United States grant number 2102953 to J.D.; ARS Project 2030-21000-056-00D to G.R.L; an USDA-NIFA Grant (2022-67013-36362) to V.K.T.; and a NERC Independent Research Fellowship (NE/T011025/1) to L.T.D.

## Author Contributions

A.G.-M., C.G., E.C.-G., S.M.R., A.R., N.C., A.B.P., C.-Y.H., K.D.L., W.J.R., F.Y.N., S.Grewal, G.S.B. and B.B.H.W. configured *Ae. tauschii* diversity panel and distributed seed. A.G.-M., A.P., A.B.P. and C.-Y.H. extracted *Ae. tauschii* DNA. S.H., M.Abbasi, M.L., N.K. and G.S.B. sampled tissue and extracted RNA from *Ae tauschii*. A.G.-M., S.H., A.F.E., L.M., A.Sharon, A.H., J.C.R., M.K., M.M., N.S., D.P., S.L., J.-Y.L., P.C., S.K., P.Z., R.F.P., D.-C.L., W.L., J.D., V.K.T., S.Grewal, C.P., H.S.C., J.E., L.T.D., S.X., Y.G., X.X., G.S.B. and B.B.H.W. outlined *Ae. tauschii* sequencing strategy and acquired DNA and RNA sequences. E.C.-G. and N.A. extracted DNA for bread wheat landraces. A.G.-M., E.C.-G., S.H., L.F.R., H.S.C. and M.Abrouk undertook sequence data quality control, curation, back-up and/or distribution. A.Salhi, X.G. and S.B. assembled *k*-mer matrix. E.C.-G. performed variant calling and filtering. A.G.-M. and E.C.-G. performed redundancy analysis. E.C.-G. and A.G.-M. performed genome-wide phylogenetic analysis and E.C.-G. performed structure analysis. A.G.-M. and H.S.C. assembled *Ae. tauschii* genomes and A.G.-M. assessed their quality. S.H. and M.Abrouk performed genome annotations. A.G.-M. performed structural variant analysis. A.G.-M. performed *k*-mer GWAS. N.A., M.P., S.Ghosh, R.C., Y.W., E.L.O, P.Z., W.B.M, W.H.P.B., B.J.S., J.A.K. and O.M. performed rust phenotyping. A.G.-M. and E.C.-G. performed *Sr33* and *Sr66* comparative genomics analyses. N.A., E.S.L, Y.D. and S.K.P. generated biparental population and mapped *Lr39*. N.A. and Y.W. identified *Lr39* candidate gene, performed VIGS, and annotated *Lr39*. E.C.-G. developed ‘Missing Link Finder’ pipeline, identified L3-enriched bread wheat lines, and assembled CWI 86942 genome. C.U. and J.Q.-C. provided prepublication access to and helped implement IBSpy pipeline, and E.C.-G., C.U. and J.Q.-C. identified L3 haploblocks in wheat and calculated their accumulative size. E.C.-G. assigned extant wheat genome segments to ancestral *Ae. tauschii* populations and determined the minimum number of hybridizations. G.Y. and H.I.A. provided scientific advice. P.K.S. and G.R.L. implemented the *Ae. tauschii* pangenome database. E.C.-G., A.G.-M., N.A., S.H., A.Salhi, M.Abrouk, S.B., G.S.B., B.B.H.W. and S.G.K. conceived and designed experiments. E.C.-G., A.G.-M., N.A., C.G., B.B.H.W. and S.G.K. designed figures. S.G.K., E.C.-G., A.G.-M and B.B.H.W. drafted first version of manuscript with inputs and edits from N.A., S.H., A.Salhi, C.G., G.S.B. and S.Ghosh. A.G.-M. and B.B.H.W. managed the Open Wild Wheat Consortium (www.openwildwheat.org).

All authors read and approved the manuscript. Authors are grouped by institution acronym and then by first name in the author list and author contribution statement – except for the first seven and last six authors.

## Competing interests

The authors declare no competing interests.

## Methods

### The *Aegilops tauschii* pangenome

#### Plant material

We compiled a database comprising 1,124 *Ae. tauschii* accessions with associated passport data in Supplementary Table 1 (Supplementary Note 1). Duplicate germplasm bank IDs were identified and passport data collated using the genesys-pgr.org database or other sources as indicated in Supplementary Table 1. From this database, seed of 228 non-redundant accessions were obtained from the Open Wild Wheat Consortium *Ae. tauschii* Diversity Panel collection deposited at the Germplasm Resource Unit (GRU) of the John Innes Centre; 48 accessions from the Cereal Crop Wild Relatives (*Triticeae*) collection of the GRU; 19 accessions from the DFW Wheat Academic Toolkit collection of the GRU that have been used as synthetic hexaploid wheat D-genome donors; 223 accessions from the Wheat Genetics Resource Center (WGRC) of Kansas State University; 34 accessions from the Plant Gene Resources of Canada (PGRC); 84 accessions donated by the Institute of Botany, Plant Physiology and Genetics of the Tajikistan National Academy of Sciences; 20 accessions donated by the Azerbaijan National Academy of Sciences; and 37 accessions donated by Quaid-i-Azam University. Accession P-99.95-1.1 was obtained from the Deposited Published Research Material collection of the GRU.

We also resequenced and analysed 60 hexaploid wheat landraces. The list can be found in Supplementary Table 15. Out of the 60 wheat landraces 57 were received from International Maize and Wheat Improvement Center (CIMMYT) and three from International Center for Agricultural Research in the Dry Areas (ICARDA).

#### Resequencing of the Ae. tauschii and hexaploid wheat accessions

In this study, we generated short-read whole genome sequencing data for 350 *Ae. tauschii* accessions (Supplementary Table 1) and 59 hexaploid wheat accessions (Supplementary Table 15). We isolated DNA following the CTAB protocol described by Abrouk *et al.*, (2020)^46^ from leaf tissue of two-week-old seedlings under prior dark treatment for 48 hours. DNA was quantified using the Qubit dsDNA HS Assay (Thermo Fisher Scientific) and purity was determined according to 260/280 and 260/230 absorbance ratios using a Nanodrop spectrophotometer. PCR-free paired-end libraries were constructed and sequenced on an Illumina Novaseq 6000 instrument, yielding a median 8.3-fold coverage per sample (ranging from 5.87 to 16.86-fold) for the *Ae. tauschii* samples and a minimum 10-fold coverage for the bread wheat samples (Supplementary Table 1, Supplementary Table 15). Library preparation and sequencing was performed as a service by Novogene.

#### Library construction and RNA sequencing

Seedlings of *Ae. tauschii* accessions TA10171, TA1675 and TA2576 were raised as 5-6 seeds per pot (6 × 6 × 10 cm) in a growth chamber at 22-24°C under long-day photoperiods of 16/8 hours day / night cycle with high-output white-fluorescent tubes until the third leaf stage (about 2-3 weeks old), and then transferred to a 4°C growth chamber with a long-day photoperiod for vernalisation. After a 9-week vernalisation period, all the plants were moved back to the original growth chamber under the controlled conditions mentioned above. In total, 45 tissue samples were collected: From each of the three accessions, three biological replicates were taken from each of: young leaf, root, stem, flag leaf and inflorescence. Samples were collected at the same time of day at approximately 5-6 hours after daylight. The seedling leaves and roots were harvested after two weeks of recovery in the original growth chamber and rinsed with water to remove soil particles. When the plants had 4-5 tillers, the stems, flag leaves and inflorescences were harvested together. The green inflorescences were collected immediately after pollination. The 5-cm-long stem sections and youngest flag leaves were measured from the top of the same inflorescences. Samples were placed in liquid nitrogen after harvest and stored at -80°C.

The samples were ground into a fine powder in liquid nitrogen in a ceramic mortar and pestle to isolate RNA using the Qiagen RNeasy Mini Kit (USA) following the manufacturer’s protocol. The quality of RNA was determined on a 1% agarose gel, and RNA concentration was measured using a NanoPhotometer (Implen, USA) at 260 nm and 280 nm. Sample collection time and relative details are listed in Supplementary Table 20. High-quality RNA samples were delivered for RNA integrity test, poly-A mRNA enrichment, library construction and PE100 sequencing using the Illumina NovaSeq system (Génome Québec, Canada).

#### PacBio HiFi genome sequencing; primary assembly of the Ae. tauschii pangenome and CWI 86942

We selected 46 *Ae. tauschii* accessions, including 11 Lineage 1 accessions, 34 Lineage 2 accessions and one Lineage 3 accession. These pangenome accessions were selected to span the geographical range of the species (Fig. 1a) and provide a collection of phenotypes related to disease and pest resistance, abiotic tolerance and agromorphological traits of strategic interest to the Open Wild Wheat Consortium for bread wheat improvement (Supplementary Table 4). We included a higher proportion of accessions from L2 relative to L1 based on reported phylogenies showing that L2 is more genetically diverse than L1^5,14,19^. A single L3 accession was selected based on low genetic diversity observed among five non-redundant L3 accessions in the phylogeny reported by Gaurav *et al.* (2022)^14^. Several accessions were selected to maximise the genetic diversity based on a core subset sampling analysis using Core Hunter (v3)^47^, employing the ‘average entry-to-nearest-entry’ distance measure, aiming to maximally represent the diversity of the panel of 242 non-redundant accessions published by Gaurav *et al.* (2022)^14^. The bread wheat landrace CWI 86942 was selected based on a high L3 *k*-mer content.

For the *Ae. tauschii* accessions, high molecular weight (HMW) genomic DNA was isolated from leaf tissue of three to four-week-old dark-treated seedlings. We followed the HMW DNA isolation protocol optimised by Driguez *et al.* (2021)^48^ for long-read sequencing. DNA integrity was confirmed using the FemtoPulse system (Agilent). DNA was quantified using the Qubit dsDNA HS Assay (Thermo Fisher Scientific) and purity was determined according to 260/280 and 260/230 ratios using a Nanodrop spectrophotometer. For the bread wheat accession CWI 86942, leaves from two-week old seedlings were collected from two different plants and high molecular weight DNA extraction was performed as mentioned above^48^. All the library preparation and Circular Consensus Sequencing (CCS) was performed on a PacBio Sequel II instrument, as a service by Novogene.

For *Ae. tauschii*, HiFi reads were assembled using hifiasm (v0.16.1)^49^ with parameters “-l0 -u -f38”. Sequencing coverage ranged from 18 to 47-fold depending on the accession, except for the three *Ae. tauschii* lineage reference accessions (TA10171, TA1675 and TA2576) for which the coverage was increased to 67 - 97-fold. For assembly validation and quality control, we used QUAST (v5.0.2)^50^ to calculate the assembly metrics, Merqury (v1.3)^51^ to estimate the base-call accuracy and *k*-mer completeness based on 21-mer produced from the short-read WGS data^14^ and BUSCO (v5.3.1)^17^ with the embryophyta_odb10 database to determine the completeness of each genome assembly.

For CWI 86942, we performed the primary contig-level assembly with 484.33 Gb of HiFi reads (32-fold coverage) using the LJA assembler (v0.2)^52^ with default parameters. Assembly metrics and QC were performed with QUAST (v5.0.2) and BUSCO (v5.3.1) with the embryophyta_odb10 database.

#### Chromosome conformation capture sequencing and chromosome-scale scaffolding

*In situ* Hi-C libraries were prepared for TA1675 and TA10171 from two-week-old *Ae. tauschii* plants according to the previously published protocol^53^. Libraries were quantified and sequenced (paired-end, 2 x 111 cycles) using the Illumina NovaSeq 6000 device (Illumina Inc., San Diego, CA, USA) at IPK Gatersleben^54^, yielding 316 million paired-end (PE) reads (150 bp) for TA1675 and 215 million PE reads for TA10171.

For TA2576, two-week-old, dark-treated leaf tissue samples were harvested and cross-linked with formaldehyde for library preparation and Hi-C sequencing by Phase Genomics, yielding 543 million PE reads (150 bp). For CWI 86942, two Omni-C libraries were generated and sequenced from two-week-old, dark-treated leaf tissue samples as a service by Dovetail Genomics. The total yield was 715 million PE reads (150 bp).

Scaffolding into pseudomolecule for TA10171, TA1675, TA2576 and CWI 86942 was performed from their primary assemblies and their specific Hi-C and Omni-C data, respectively. Hi-C and Omni-C reads were processed with Juicer (v1.6)^55^ (for the Hi-C reads, parameter: -s DpnII) to convert raw fastq reads to chromatin contacts and remove duplicates. The chromatin contacts were used to scaffold the contig-level assemblies using 3D-DNA (v190716)^56^ (using run-asm-pipeline.sh with -r 0 parameter). Scaffolds were visualised, manually oriented and ordered using Juicebox (v2.20.00)^57^.

#### RagTag assembly of the 43 pangenome accessions

The remaining 43 contig-level assemblies were scaffolded into chromosome-scale assemblies using RagTag (v2.1.0)^58^ and the three high-quality genomes (TA10171, TA1675 and TA2576) as anchors. Briefly, the primary contig-level assemblies were scaffolded using RagTag and the -scaffold option against the respective chromosome-scale reference assemblies generated in this study. The scaffolded assemblies were validated with dot-plots generated using MashMap (v3.0.6)^59^ against the corresponding reference assembly.

#### Repeat and gene annotation

Paired-end RNAseq reads for TA10171, TA1675 and TA2576 were first cleaned using Trimmomatic (v0.40)^60^ with the following settings “ILLUMINACLIP:TruSeq3-PE.fa:2:30:10:2:True LEADING:30 TRAILING:30 MINLEN:36”. Trimmed paired-end reads were aligned to the corresponding genome assembly using STAR (v2.7.10b)^61^ with the parameters “--twopassMode basic --outFilterMismatchNMax 5 --outFilterMatchNminOverLread 0.80 --alignMatesGapMax 100000 -- outSAMstrandField intronMotif --runMode alignReads” and the results were filtered and sorted using SAMtools (v1.10)^62^. Then, the braker (v3.0.3)^63-65^ pipeline was used to predict *de novo* gene models using RNA-Seq and protein data mode with the *Viridiplantae* protein models provided by OrthoDB (v11). Predicted gene annotations obtained from braker were processed using a combination of NCBI BLAST+ (v2.9.0-2)^66^, AGAT (v1.2.1) (https://github.com/NBISweden/AGAT), InterProScan (v5.64-96.0)^67,68^, and R (v4.2.0). Outputs from braker3 were first converted to gff3 and CDS and protein sequences were extracted using “agat_sp_extract_sequences.pl” from AGAT package. BLASTn was used to perform a reciprocal BLAST of the predicted CDS against themselves, and a unidirectional BLAST against the Ensembl nrTEplantsJune2020.fa repetitive elements database, using default search parameters. The putative functions for each annotated gene model were predicted using InterProScan with default parameters for the following databases: FunFam, SFLD, PANTHER, Gene3D, PRINTS, Coils, SUPERFAMILY, SMART, CDD, PIRSR, ProSitePatterns, AntiFam, Pfam, MobiDBLite, PIRSF, NCBIfam. R (v4.2.0) (in R studio) was used to visualise and filter these results. Predicted transcripts with fewer than 50 exact and fewer than 150 inexact self-BLAST results were retained. Predicted transcripts were retained from the final *de novo* annotation if there were (i) no exact matches to the transposable elements database and (ii) at least on domain predicted by any of: FunFam, PANTHER, Gene3D, SUPERFAMILY, ProSitePatterns, Pfam, CDD, InterPro. Predicted genes were considered as “Low Confidence” if there were no exact matches to the database of original transcript predictions. The remaining annotated genes were considered as “High Confidence”. Validation and annotation completeness was performed using agat_sp_statistics.pl and BUSCO (v5.4.7)^17^ running in transcriptome mode with the poales_odb10 database.

Repeat annotation was performed using RepeatMasker (v4.1.2-p1)^69^ and the EnsemblnrTEplantsJune2020 repetitive elements database^70^ using the RMBlast engine.

For bread wheat accession CWI 86942, gene model prediction was performed using a lifting approach similarly to the one described in Abrouk *et al.* (2023)^71^ with a combination of liftoff (v1.6.3)^72^, AGAT and gffread (v0.11.7)^73^. Briefly, gene model annotations of hexaploid wheat line Chinese Spring, Kariega, Fielder, Arina*LrFor*, Julius and Norin61 were independently transferred using liftoff (parameters: -a 0.9 -s 0.9 -copies -exclude_partial -polish) and all the output gff files were merged into a single file using the Perl script “agat_sp_merge_annotations.pl”. The merged file was then post-processed using gffread tools (parameters: --keep-genes -N -J) to retain transcripts with start and stop codons, and to discard transcripts with (i) premature stop codons and/or (ii) having introns with non-canonical splice sites. In total, 147,646 gene models were predicted for which the putative functional annotations were assigned using a protein comparison against the UniProt database (2021_03) using DIAMOND (v2.1.8)^74^ (parameters: -f 6 -k1 -e 1e-6). PFAM domain signatures and GO were assigned using InterproScan (v5.55-88.09)^67,68^. The BUSCO score showed a completeness of 99.2% (96.4% duplicated) with the poales_odb10 database^17^.

#### Cumulative k-mer content across the pangenome

We computed the *k*-mer accumulation across the 46 *Ae. tauschii* pangenome accessions by performing pairwise comparisons between short-read-derived *k*-mer datasets (*k*=51) to extract core and shell *k*-mers. First, we ordered the accessions using Core Hunter (v3)^47^, employing the ‘average entry-to-nearest-entry’ distance measure to maximise the mean distance between a given accession and the next accession in the list. In that order we then compared the *k*-mer sets between the first and second accessions to extract and count shared *k*-mers and *k*-mers unique to accessions 1 and 2 using the comm command. The shared *k*-mers from the previous comparison were then compared to the *k*-mer set of accession 3 to extract and count the cumulative shared *k*-mers and unique *k*-mers. This comparison was performed sequentially until accession 46. The cumulative shared and shell *k*-mer counts were fitted to a logarithmic function [y = a+b*log(x)] using the Python function optimize.curve_fit from SciPy library (v1.8.0)^75^. The fitted data was plotted using the Python seaborn library (v0.11.2)^76^ to visualise the *k*-mer-based pangenome accumulation curve.

#### SNP calling

Fastq raw reads were trimmed using Trimmomatic (v0.38)^60^ with the following settings “ILLUMINACLIP:adapters.fa:2:30:10 LEADING:3 TRAILING:3 SLIDINGWINDOW:4:15 MINLEN:36”. Cleaned reads were mapped on the TA1675 assembly using BWA mem (v0.7.17)^77^ and sorted with SAMtools (v1.8)^62^. Variants were called using BCFtools mpileup (v1.9)^62^ with the following setting “-C 60 -q 5 -Q 20”, and only SNPs were retained as variants. The filtering was performed using BCFtools, retaining only sites with a maximum depth of 40, a quality higher than 100 and an allele count higher than one. For quality check, we counted the percentage of divergent sites using re-sequencing data from TA1675 against the chromosome-scale TA1675 reference assembly, revealing an error rate of 0.18%. We further computed the SNP density across the chromosomes and calculated allele frequency (Extended Data Fig. 8 a-c, Supplementary Tables 21-23). In total, 957 *Ae. tauschii* and 59 wheat landraces accessions reached the quality threshold of a coverage higher than 5-fold after trimming and were used for SNP calling. The phylogenetic tree was built from the filtered SNPs using vcfkit (v0.1.6)^78^ with the UPGMA algorithm. Ancestry analysis was performed using the sNMF (Fast and Efficient Estimation of Individual Ancestry Coefficients) approach available in the LEA R package (v3.10.2)^79^. For each run, we performed 20 repetitions using the following parameters “alpha = 10, tolerance = 0.00001, iterations = 300” up to *K*=28. Supplementary Table 24 shows the minimum cross-entropy values for 20 sNMF runs across different values of *K*.

We estimated the nucleotide diversity (π) in the 493 non-redundant accession of *Ae. tauschii* using the filtered SNP calls against the TA1675 reference assembly. We calculated π over 10 kb windows of the genome using VCFtools (v0.1.16)^80^ (parameter --window-pi 10000).

#### k-mer matrix generation

We developed an optimised *k*-mer matrix workflow to generate a presence/absence *k*-mer matrix for large diversity panels (Supplementary Note 2) (https://github.com/githubcbrc/KGWASMatrix). We counted *k*-mers (*k* = 51) in raw sequencing data for 350 accessions generated in this study, 306 accessions published by Gaurav *et al.* (2022)^14^, 275 accessions published by Zhou *et al.* (2021)^15^ and 24 accessions by Zhao *et al.* (2023)^6^. The 35 accessions with less than 5-fold sequencing coverage were discarded to avoid affecting the *k*-mer count. *k-*mers were filtered by a minimum occurrence of six across accessions and a maximum occurrence of (*N*-6), being *N* the total number of accessions.

#### Redundancy analysis

A redundancy analysis was performed using a subset of 100,000 random *k*-mers sampled from the *k*-mer matrix of 920 *Ae. tauschii* accessions. The complete matrix contained 10,078,115,665 *k*-mers. The pairwise comparisons between accessions were performed by computing the sum of the presence-absence values (0 and 1) per *k*-mer between two accessions of the matrix. To determine the divergence, we computed the total number of 1 present in the summed string, each one corresponding to a difference in the presence/absence of the *k*-mer in the two compared accessions. In the sum, the presence of a *k*-mer in two accessions would result in a 2 and the absence in both accessions in a 0. A threshold of 96% shared *k*-mers was used to call redundancy based on control lines determined by Gaurav *et al.* (2022)^14^ to be genetically redundant based on a SNP analysis.

#### Structural variant (SV) calling

We determined the structural variation across the pangenome with reference to the chromosome-scale assembly of TA1675. Structural variants (SVs) of >50 bp in length were called using the PacBio structural variant calling and analysis suite (pbsv) (v2.9.0) and following the pipeline described at (https://github.com/PacificBiosciences/pbsv). Briefly, HiFi sequencing reads in bam format were aligned to the reference genome using pbmm2 aligner (v1.10.0) (https://github.com/PacificBiosciences/pbmm2). The bam file was indexed as CSI suitable for larger genomes. Signatures of structural variation were detected and structural variants were called per accession in vcf format, then concatenated into a single bed file per lineage.

### The *Ae. tauschii* pangenome facilitates gene discovery

#### k-mer-based genome-wide association in Aegilops tauschii

We followed the *k*-mer GWAS (*k*GWAS) pipeline described by Gaurav *et al.* (2022)^14^ using the Python scripts available at (https://github.com/wheatgenetics/owwc/tree/master/kGWAS) and the phenotype data for stem rust and leaf rust available for this panel to specifically run the association mapping and plotting using default parameters. The association mapping analyses showing the effect of assembly quality in *Ae. tauschii* accession TA1662 were performed using previously published phenotype data for reaction to *Puccinia graminis* f. sp. *tritici* (*Pgt*) race QTHJC^14^. The *k*GWAS for leaf rust to identify *Lr39* in *Ae. tauschii* accession TA1675 was performed using phenotype data for reaction to *Puccinia triticina* (*Pt*) race BBBDB (Supplementary Table 25).

#### SrTA1662 haplotype analysis

To identify the *SrTA1662* locus in the contig-level assembly of *Ae. tauschii* accession TA1662, we performed a BLASTn (v2.12.0)^66^ search of the *SrTA1662* gene sequence (GenBank ID MW526949.1) published by Gaurav *et al.* (2022)^14^. To identify the *SR33* locus in the contig-level assembly of *Ae. tauschii* accession CPI 110799 (the original source of *Sr33*), we searched for the *RGA1e* (*Sr33*) gene sequence (GenBank ID KF031291.1) published by Periyannan *et al.* (2013)^22^. *RGA1e* gene sequences were also searched against the contig-level assemblies of accessions AUS 18911 (KF031299.1) and AUS 18913 (KF031284.1)^22^. For all accessions, the genes were found within a single contig that located to the chromosome arm 1DS based on the scaffolding against the TA1675 reference assembly. In the four accessions, we performed BLASTn searches for additional *Ae. tauschii* Resistance Gene Analogues (*RGA1a-d*, *RGA2a-b*, *RGA3a*) reported by Periyannan *et al.* (2013)^22^ (GenBank ID KF031285.1-KF031299.1). To confirm that this region is orthologues to the *Mla* locus in barley, we searched for the presence of the pumilio (*Bpm*) gene homologue and subtilisin-chymotrypsin inhibitor (*CI*) genes in gene-lifting annotations for AUS 18911, AUS 18913, CPI 110799 and TA1662. *Bpm* and *CI* genes were previously reported flanking Resistance Gene Homologs (*RGH*) of the *Mla* locus^81^. The gene lifting annotations were generated using litftoff v1.6.1^72^ with default parameters based on the TA1675 genome annotation.

#### Phylogenetic analysis of Resistance Gene Analogues (RGA) in Aegilops tauschii

To provide further evidence for the homology of the *SrTA1662* (*SR66*) and *SR33* loci in *Ae. tauschii* and the *Mla* locus in barley, we performed phylogenetic analyses of *RGA* and *RGH* gene sequences. Clustal algorithm with default parameters was used for the DNA sequence multiple alignment. We used the unweighted UPGMA algorithm with bootstrap testing to support the tree topology with 5000 replicates. The phylogenetic analyses were performed using MEGA (v11)^82,83^.

#### Leaf rust inoculations and association studies

The evaluation of resistance and susceptibility in 149 *Ae. tauschii* L2 accessions was conducted against the North American *Pt* race isolate BBBDB 1-1^84^ using seedlings organised in cone racks. Every cone rack housed 98 cones, and each cone was sown with three seeds. The primary leaves of seedlings, aged 8-9 days, were subjected to inoculation by distributing 1 mL of inoculum per cone rack, which consisted of 15 mg of spores in 1 mL of Soltrol 170 (Chevron Phillips Chemical Company, USA). The delivery to each plant was 0.05 mg of urediniospores. Post-inoculation, the phytotoxicity from the oil carrier, Soltrol 170, was mitigated by mildly fanning the leaves for two hours under the illumination of 400-watt HPS bulbs to expedite the evaporation of the carrier oil. The seedlings, once inoculated, were placed in mist chambers maintained at 22°C, where 100% humidity was sustained using a domestic ultrasonic humidifier for a period of 16-18 h in the absence of light. Subsequently, the seedlings were transferred to a greenhouse with a 16-hour day cycle, maintaining nocturnal and diurnal temperatures at 15°C and 20°C, respectively. The phenotypic assessment of disease was undertaken at 10 and 12 dpi using an infection type scoring range of 0 to 3+4, as standardised by Long and Kolmer^85^, and depicted as the mean of three individual replicates per accession (Supplementary Table 25).

For use in GWAS, the qualitative scores were converted to a quantitative score by assigning numerical values to the infection types. This was achieved by the *k*GWAS pipeline (https://github.com/wheatgenetics/owwc/tree/master/kGWAS) that performs Stakman IT to numeric scale 1 conversion (RunAssociation_GLM.py with -st parameter).

#### Bi-parental mapping of LR39 and candidate gene identification

An *Ae. tauschii* bi-parental mapping population (*n* = 123) was generated by crossing the leaf rust resistant *Ae. tauschii* accession CPI 110672 (synonymous TA1675) with a leaf rust susceptible accession CPI 110717. The mapping population was segregating for a single dominant leaf rust resistance gene (*P* = 0.606) when inoculated with the Australian *Pt* isolate 26-1,3 (PBI culture no 316) phenotyped at the Plant Breeding Institute, Cobbitty^28^. Bulk segregant analysis of selected homozygous resistant and susceptible F_2_ progenies with the 90K iSelect single nucleotide polymorphism (SNP) array^86^ placed the leaf rust resistance locus to chromosome 2DS. The mapping population was further genotyped with markers derived from the 90K iSelect SNP array, the TA1675 genomic sequence, and a marker closely linked to *LR39* (*Xgdm35*) (Supplementary Table 26)^26,87^. Linkage analysis was performed using MapDisto (v2.0)^88^ with default parameters such as LOD (logarithm of the odds) threshold of 3.0, maximum recombination frequency of 0.3 and removal of loci with 10% missing data. Genetic distances were calculated using the Kosambi mapping function, and the map was created using MapChart (v2.32)^89^. Markers flanking *LR39* were anchored to the TA1675 reference assembly. Annotated high-confidence genes at the delimited physical interval were screened for protein homology using BLASTp to identify diversity between TA1675 and AL8/78 (Supplementary Table 12). The conserved domain and critical residues of WTK and NLR were identified using the amino acid sequences in the NCBI Conserved Domain search (https://www.ncbi.nlm.nih.gov/Structure/cdd/wrpsb.cgi) and InterPro (https://www.ebi.ac.uk/interpro/search/sequence/) databases. The polymorphic SNP corresponding to R457I in WTK was converted to *Lr39* diagnostic KASP marker (Supplementary Table 27).

#### Virus-induced gene silencing (VIGS)

To develop candidate gene-specific VIGS probes, the predicted coding sequences of candidate genes were searched against the TA1675 transcriptome database using siRNA-Finder (siFi21) software (v1.2.3-0008)^90^. Based on the RNA interference (RNAi) design plot, regions predicted to have a higher number of efficient probes and fewer off-targets were used for designing silencing probes for the WTK (LrSi2:258 bp, LrSi6:254 bp, LrSi7:248 bp) and the NLR (LrSi3:234 bp, LrSi8:257 bp) candidate genes. The silencing probe sequences were verified for specificity using a BLAST search against the TA1675 reference assembly (<80% sequence identity for hits other than the target candidate). Designed probes were flanked by *Xba*I and *Apa*I and synthesised at GenScript Biotech followed by cloning into the BSMV-γ vector in an antisense direction. The resulting constructs were transformed into *Agrobacterium tumefaciens* strain GV3101. The *Agrobacterium* clones were grown overnight at 28°C in lysogeny broth with appropriate antibiotics. Cells were collected by centrifugation at 3,500 *g* for 10 min, then re-suspended using infiltration buffer (10mM MgCl_2_, 10 mM MES pH 6.5 with KOH buffer and 150 mM Acetosyringone) and adjusted to an OD_600_ of 1.0 followed by incubation at 28°C for 3 h. Equal volumes of BSMV-α and BSMV-β were mixed with respective BSMV-γ silencing probes or BSMV-γGFP and infiltrated into *Nicotiana benthamiana* leaves. Infiltrated leaves were collected 5 days after infiltration and homogenised with virus inoculation buffer (10 mM monopotassium phosphate containing 1% Celite (Thermo Fisher Scientific, 68855-54-9)). The homogenate containing viral particles was rub inoculated onto five to ten seedlings of TA1675 at the three leaf stage. After two weeks of recovery and viral symptoms appearing, the seedlings were inoculated with *Pt* isolate B9414. Prior to inoculating TA1675, *Pt* isolate B9414 was propagated on seedlings of the susceptible wheat cultivar Thatcher. Freshly collected urediniospores were suspended in IsoparL and sprayed onto plants using a high-pressure air sprayer. After inoculation, plants were placed in the dark overnight in an incubation box equipped with a humidifier and then transferred to a growth chamber with a 16/8 h day/night cycle, with 21°C/18°C growth conditions. Leaf rust phenotypes were recorded at 12 days after inoculation by scanning the leaves at 600 dots per inch on an Epson Perfection V850 Pro scanner.

#### PCR conditions

A 20 μl PCR containing 100 ng of genomic DNA, 1X GoTaq Flexi green buffer, 1.5mM MgCl2, 200 μM dNTP, 200 nM primers and one unit of Taq polymerase (M829B, Promega, USA) was used for various fragment amplifications. Primer sequences are shown in Supplementary Table 27. A touchdown PCR protocol was used as follows: initial denaturation at 94°C for 30 s; annealing at 65°C for 30 s, decreasing by 1°C per cycle; and extension at 72°C for 60 s, followed by repeating these steps for 14 cycles. After enrichment, the program continued for 29 cycles as follows: 94°C for 30 s, 58°C for 30 s, and 72°C for 60 s. PCR products of cleaved amplified polymorphic sequence (CAPS) markers were digested with appropriate restriction enzymes by following the manufacturer’s instructions. A 5 μl reaction (2.5 μl of KASP Master Mix (Low ROX KBS-1016-016), 0.07 μl of assay mix and 2.5 μl (25 ng) of DNA) was used for KASP markers. PCR cycling was performed in an ABI QuantStudio 6 Flex Real-Time PCR machine as follows: preread at 30°C for 60 s; hold stage at 94°C for 15 min; and then ten touchdown cycles (94°C for 20 s; touchdown at 61°C, decreasing by 0.6°C per cycle for 60 s), followed by 29 additional cycles (94°C for 20 s; 55°C for 60 s). The plates were then read at 30°C for endpoint fluorescent measurement.

### Tracing lineage-specific *Ae. tauschii* haplotype blocks in the bread wheat genome

#### Missing link finder pipeline

We generated canonical 51-mers for each of the 82,293 genotyped wheat accessions from Sansaloni *et al*. (2020)^32^ using their respective DArTseq markers and Jellyfish (v 2.3.0)^91^. For each accession, the *k*-mers were sorted and stored as text files. From the *k-*mer matrix available from Gaurav *et al.* (2022)^14^, *k*-mers present only in *Ae. tauschii* Lineage 3 (L3) were extracted, sorted, and stored as text files. Pairwise comparisons of the sample-specific *k*-mers from the 82,293 wheat accessions and the L3-specific *k*-mers were performed using the “comm” bash command. Jaccard indices were computed with the following formula, where A is the set of *k*-mers from a single accession and L is the L3-specific *k*-mer set.

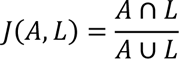

The script is available on the following Github page (https://github.com/emilecg/wheat_evolution).

#### Determining the extent of Lineage 3 in wheat lines using whole genome resequencing data of 59 landraces

Raw reads were trimmed using Trimmomatic (v0.38)^60^ with the following settings “ILLUMINACLIP:adapters.fa:2:30:10 LEADING:3 TRAILING:3 SLIDINGWINDOW:4:15 MINLEN:36”. KMC (v3.1.2)^92^ was used to generate 31-mer sets for the 59 resequenced wheat landraces (Supplementary Table 15). IBSpy (v0.4.6)^13^ was run with TA2576 as a reference and the bread wheat landraces as queries with a *k*-mer size of 31 and a window size of 50,000 bp as parameters. A variation score threshold of ≤150 was used to determine how many windows were in common between the L3 reference and the landraces. IBSpy variation values of ≤150 were determined to be optimal to account for the relatively low inter-lineage *variation* present in the Lineage 3 (Extended Data Fig. 9, Supplementary Table 28). The percentage of matching 50-kb windows was used as a proxy to determine the extent of introgression in the landraces.

#### Differences in genes in the 135-Mb L3 introgression block on chromosome 1D

The protein sequences of genes annotated in the interval of the introgression on the Kariega genome and on the CWI 86942 genome were compared using DIAMOND and visualised with the Persephone^®^ genome browser. In case genes were annotated in both the genomes, their amino acid sequences were aligned using the Needleman-Wunsch algorithm to determine the percentage of identity. The absence of genes in one of the two annotations was investigated manually with the BLAST algorithm integrated into Persephone^®^. Annotated genes found to be part of transposable elements were excluded from the analysis.

#### Presence of the 135-Mb L3 haplotype block on chromosome 1D in wheat landraces

The presence of the 135-Mb L3 haplotype block was manually confirmed in 12 out of the 126 wheat landraces (Supplementary Table 15). CWI 86942 and another Georgian landrace (CWI 86929) had the largest block size (Extended Data Fig. 6d).

To further determine how widespread the presence of the chromosome 1D L3 segment was, we downloaded the IBSpy variation file from 1,035 hexaploid wheat accessions (827 landraces and 208 modern cultivars) and L3 line BW_01028 (https://opendata.earlham.ac.uk/wheat/under_license/toronto/WatSeq_2023-09-15_landrace_modern_Variation_Data/IBSpy_variations_10WheatGenomes/) against the Chinese Spring RefSeq v1.0 assembly^35^. We found an additional 20 wheat accessions that carry at least parts of this segment. We defined the start and end of the L3 segments in these 20 accessions by determining the difference between the variation value of BW_01028 (L3) and the corresponding variation value of the twenty accessions. If the difference was ≤150, we defined the accession to carry the L3 segment.

#### Bread wheat D genome subpopulations contribution

The approach used for a quantitative estimation of the contributions of the different subpopulations to the D genomes is described in Supplementary Note 3. The manual curation process that allowed counting the minimal number of hybridizations required to explain the presence of different haplotypes is described in Supplementary Note 4.

#### Data visualisation

We used Circos (v0.69-8)^93^ to generate the circos plot in Fig.1c, including the tracks showing gene density per 10 Mb windows for TA1675, repeat density per 10 Mb windows for TA1675, nucleotide diversity across the 493 non-redundant accessions of *Ae. tauschii* based on variant calls (SNPs) per 10 kb window relative TA1675, and structural variant (SV) density across 10 kb windows in L1, L2 and L3 accessions of the pangenome. We used the R package karyoploteR (v1.20.3)^94^ for the haplotype representation of the chromosomes in figures 3d, 3e, 4b and Extended Data Fig. 7a. The remaining plots are produced with ggplot2 (v3.4.2)^95^ and the Python seaborn library (v0.11.2)^76^. Maps in figures: 1a, 3c, 4a and Extended Data Fig. 7b were generated using QGIS (v3.32.3)

## Data availability

The *Ae. tauschii* genomic resources are made available under the Toronto agreement. Data utilization can be requested through the Open Wild Wheat Consortium (https://openwildwheat.org/aegilops-tauschii/).

## Notes

### Competing Interest Statement

The authors have declared no competing interest.

