## Extended data for "Origin and evolution of the bread wheat D genome"

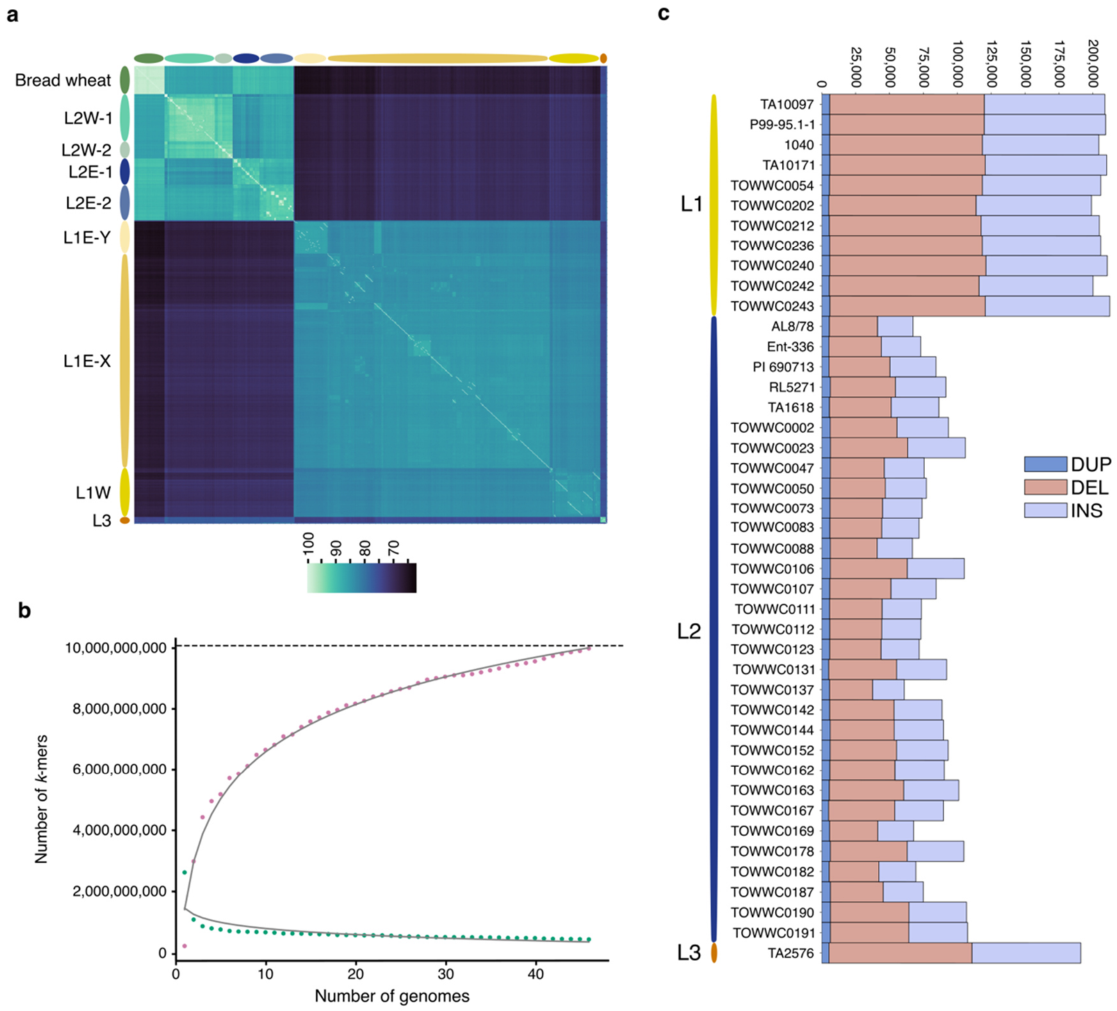


**Extended Data Fig. 1. The *Aegilops tauschii* pangenome. a**, Clustered heatmap showing SNP-based pairwise identity across 957 *Ae. tauschii* accessions and 59 bread wheat landraces. The different *Ae. tauschii* subpopulations are indicated on the left. **b**, Accumulation curves for core *k*-mers (bottom curve) and shell *k*-mers (top curve) across the 46 *Ae. tauschii* pangenome accessions. The untransformed data points are shown in pink (shell *k*-mers) and green (core *k*-mers). The dashed black line indicates the total number of *k*-mers in the diversity panel. **c**, Number of structural variants across the *Ae. tauschii* pangenome accessions relative to the chromosome-scale assembly of L2 accession TA1675. Shown are duplications (DUP), deletions (DEL), and insertions (INS) ranging from 50 bp to 100 kb.

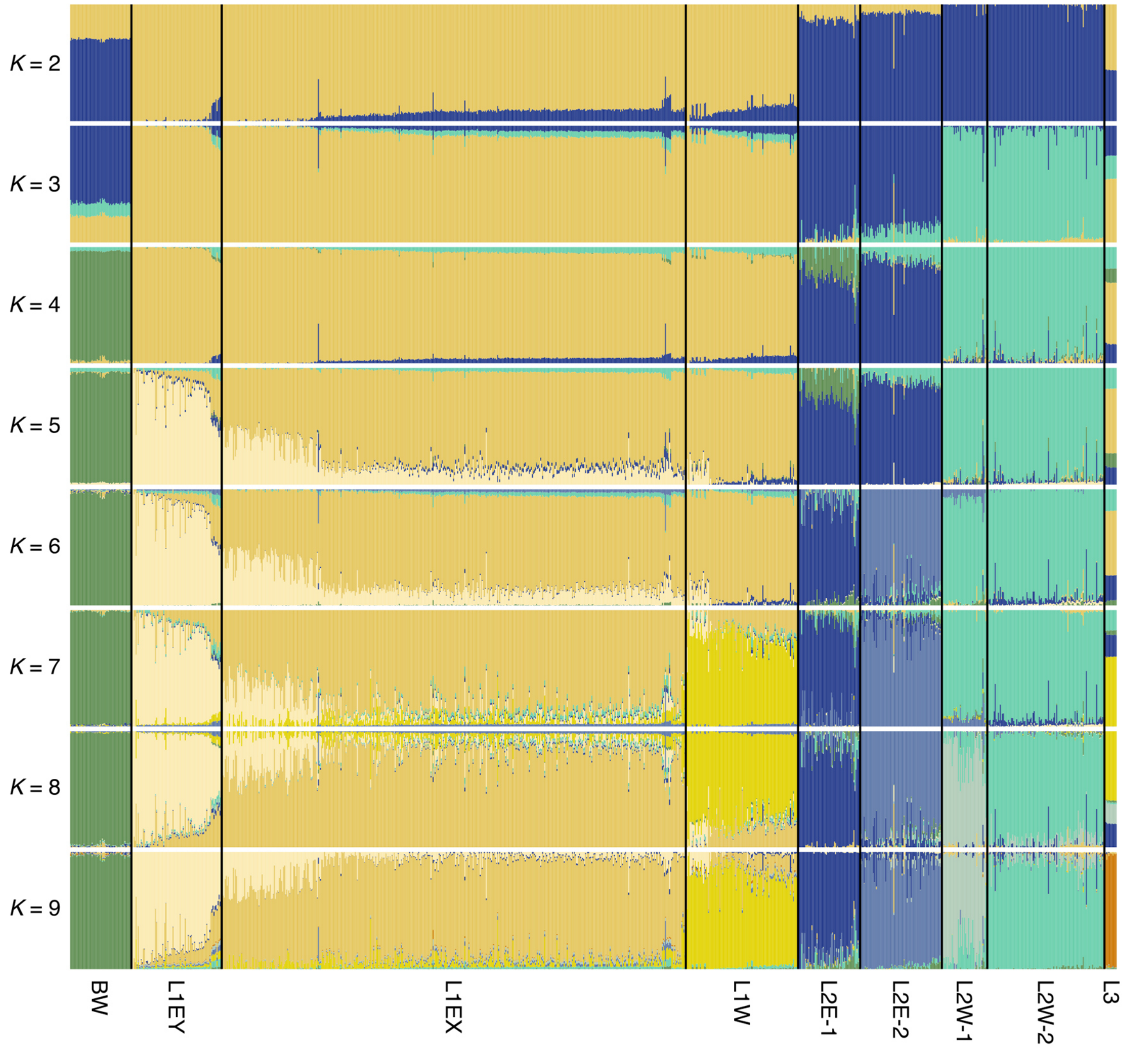


**Extended Data Fig. 2.** ***Ae. tauschii*** **population structure from** ***K*****=2 to** ***K*****=9.** Each vertical bar represents an accession and the bars are filled by colors representing the proportion of each ancestry. The subpopulation designations are described in the main text. BW = bread wheat.

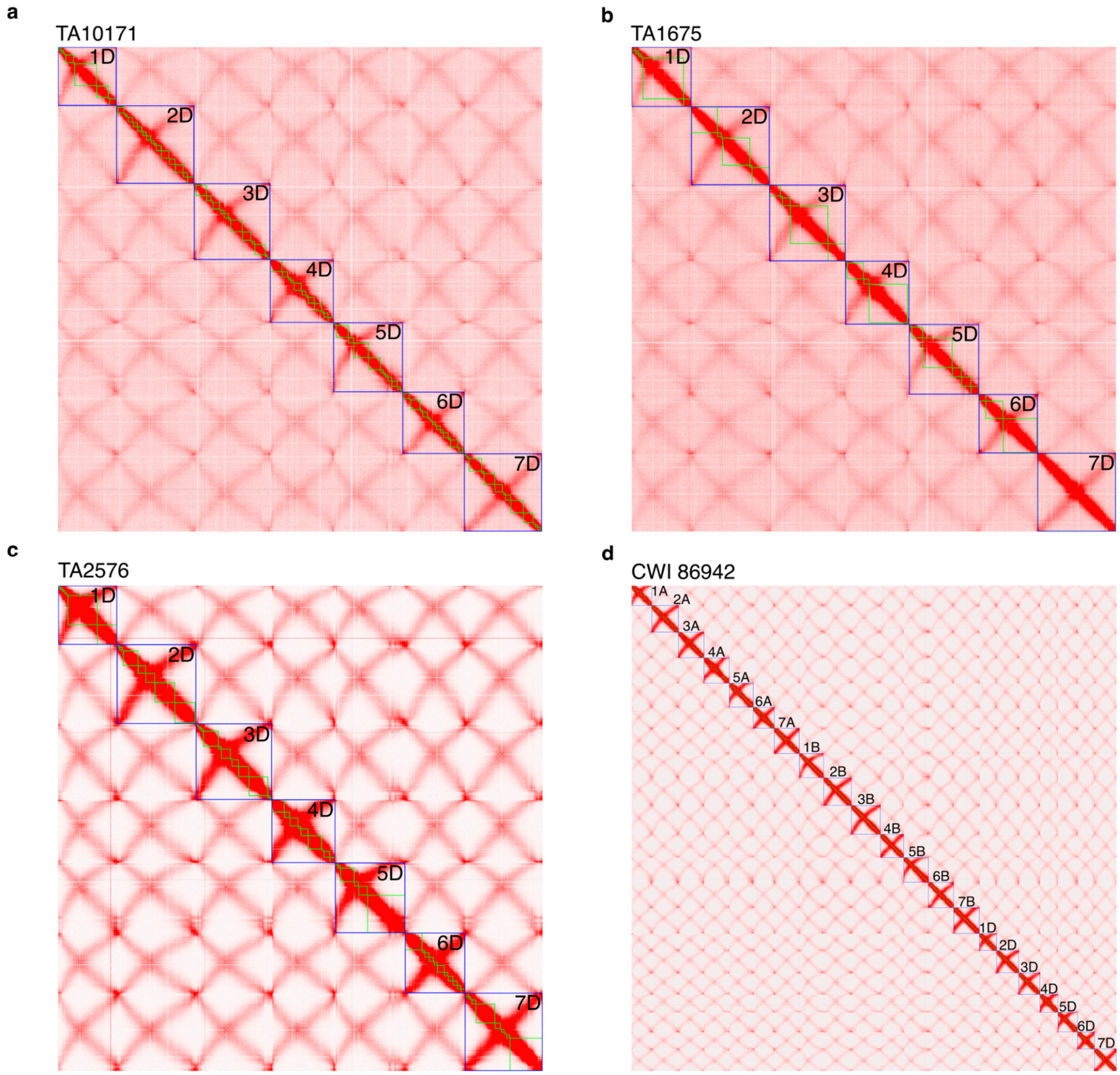


**Extended Data Fig. 3. Chromosome contact maps of *Ae. tauschii* accessions TA10171 (a), TA1675 (b), TA2576 (c), and bread wheat accession CWI 86942 (d).** Green boxes represent individual PacBio contigs. Blue boxes indicate chromosomes. Chromosome 7D of TA1675 was assembled as a single PacBio contig.

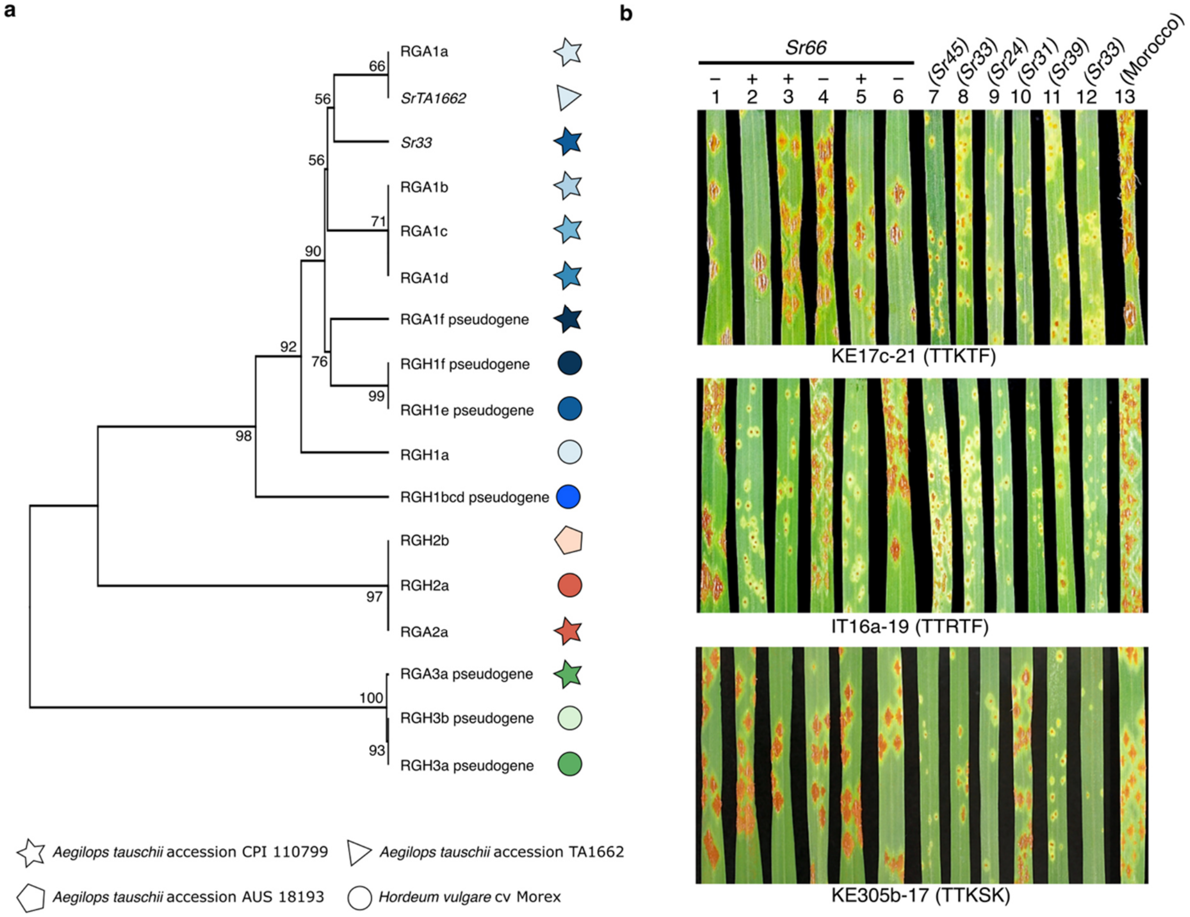


**Extended Data Fig. 4. Haplotype analysis leads to the designation of stem rust resistance gene *Sr66*.** **a**, Phylogeny showing the relationship across *Mla* genes from *Ae. tauschii* and barley. Resistance Gene Analogs (RGA) represent *Ae. tauschii* and Resistance Gene Homologs (RGH) represent barley cultivar Morex. The *Ae. tauschii* RGA gene sequences were derived from different accessions (Supplementary Table 9). RGA/RGH families 1, 2 and 3 are indicated in blue, red and green, respectively. The tree was constructed using the unweighted UPGMA algorithm. Bootstrap support values are shown based on 5,000 replicates. **b**, *SrTA1662* (*Sr66*) and *Sr33* display different race specificities. Reactions to *Puccinia graminis* f. sp. *tritici* isolates KE17c-21 (race TTKTF), IT16a-19 (TTRTF), and KE305b-17 (TTKSK) of transgenic *SrTA1662* (*Sr66*) wheat lines and non-transgenic nulls (1 to 6) and wheat *Sr* gene introgression lines and controls (7 to 13). 1, Fielder null (DPRM0050); 2, *Sr66* (DPRM0051); 3, *Sr66* (DPRM0059); 4, Fielder null (DPRM0062); 5, *Sr66* (DPRM0071); 6, Fielder null (DPRM0072); 7. *Sr45* (RL5406); 8. *Sr33* (RL5405); 9. *Sr24* (LcSr24Ag); 10. *Sr31* (Little Club/Agent (CI 13523)); 11. *Sr39* (RL5711); 12. *Sr33* (Chinese Spring); 13. cv. Morocco.

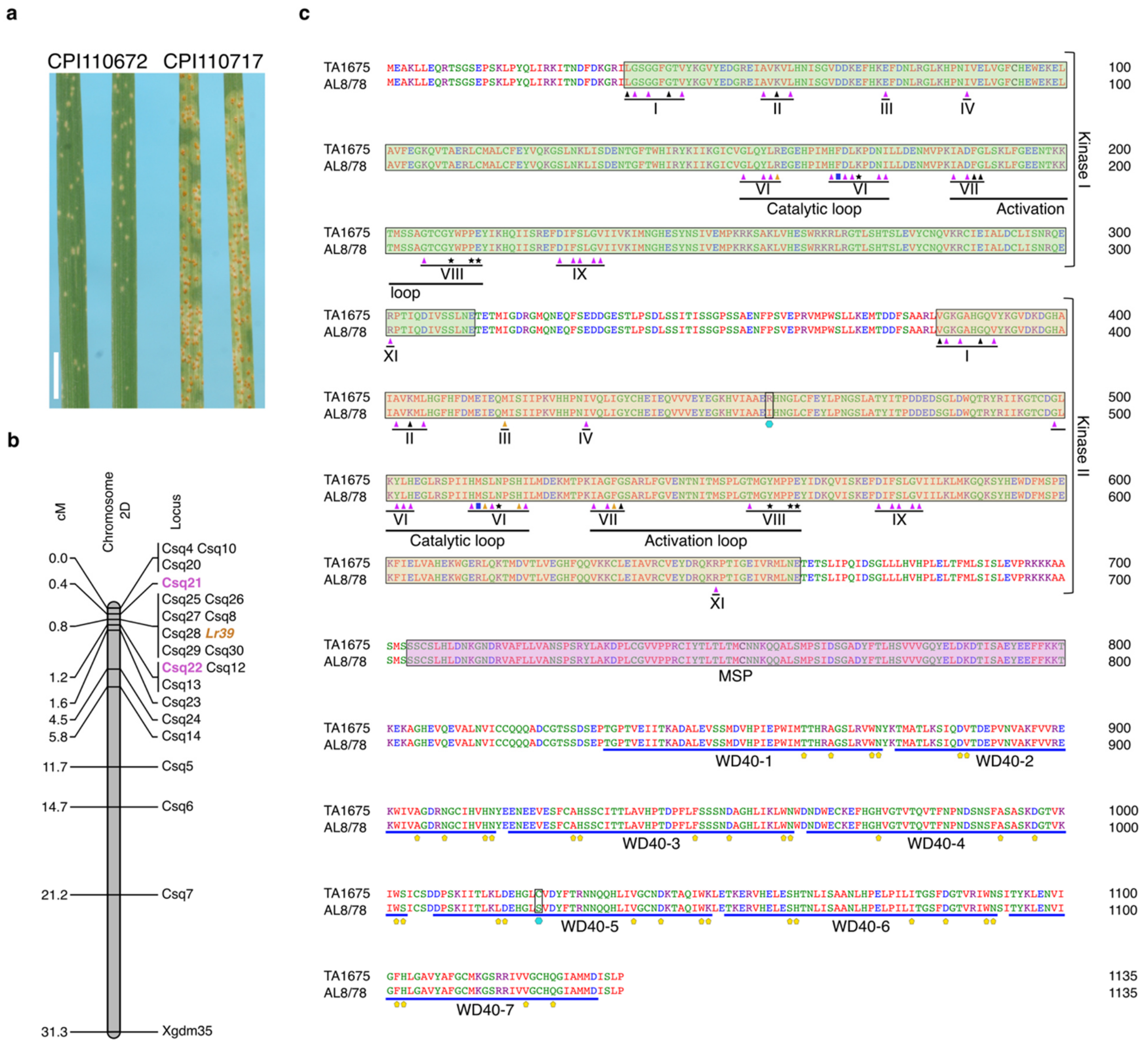


**Extended Data Fig. 5. Bi-parental genetic mapping of *LR39* and analysis of key conserved domains in Lr39. a**, Phenotypes of *Ae. tauschii* parents inoculated with the *Puccinia triticina* race Pt 26-1,3 (accession 316). CPI110672 (synonymous TA1675) carries *Lr39*. CPI110717 is the susceptible parent. Scale bar = 1 cm. **b**, Fine mapping of *LR39* in chromosome arm 2DS. Markers *Csq21* and *Csq22* are flanking the *LR39* locus whereas *Csq8*, *Csq25*, *Csq26*, *Csq27*, *Csq28*, *Csq29* and *Csq30* are co-segregating. **c**, Analysis of key conserved domains of the Lr39 protein. The kinase 1 domain is highlighted by a green box, kinase 2 by a yellow box, the major sperm protein (MSP) domain by a pink box, and the seven WD40-repeats are underlined by blue lines. Roman numerals represent conserved kinase subdomains. Black triangles = ATP binding site predicted by InterPro; Magenta triangles = key conserved residues, black asterisks = putative substrate binding site, blue squares = residue determining RD and non-RD kinases, brown triangles = polymorphism in the key conserved residues. In kinase 1, a key residue histidine is replaced by arginine in subdomain VI. In kinase 2, substitutions of residues glutamic acid to methionine in subdomain III, aspartic acid to serine and asparagine to histidine in subdomain VI form a catalytic loop, and aspartic acid to glycine in subdomain VII in the activation loop. Yellow pentagons = key conserved residues of WD40 repeats predicted by InterPro. Cyan hexagon = two polymorphic residues of TA1675 compared to AL8/78.

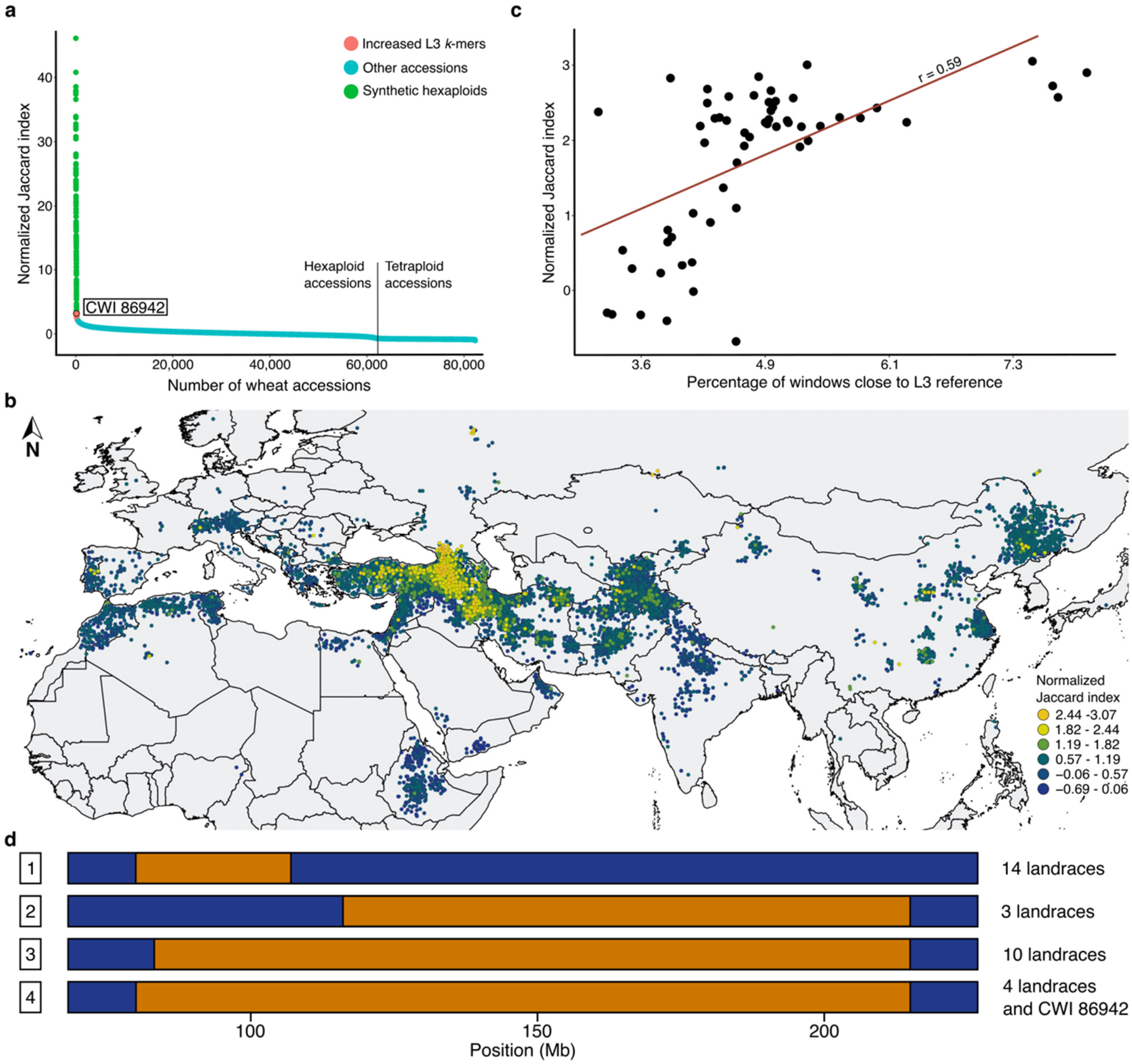


**Extended Data Fig. 6. Tracing lineage-specific *Ae. tauschii* haplotype blocks in bread wheat**. **a**, Normalized Jaccard scores across 82,293 wheat accessions (including the 139 synthetic hexaploid wheats). Green indicates 139 synthetic hexaploid wheat accessions with *k*-mer enrichments of up to 40-fold. Red indicates bread wheat landraces with increased (2 to 3-fold) normalized Jaccard index. **b**, The Jaccard indices show a gradual decline with increasing geographic distance from Georgia. Dots represent individual bread wheat accessions for which exact coordinates were available. Colors represent different normalized Jaccard indices. **c**, Correlation between normalized Jaccard indices and the percentage of L3 genome based on whole-genome sequencing data. **d**, Diagram of a portion of chromosome arm 1DS. The chromosome positions indicated in Mb are according to the CWI 86942 assembly. Haplotype blocks corresponding to *Ae. tauschii* L2 are indicated in blue, and L3 in orange. Shown are different lengths of the L3 haplotype segment in various bread wheat lines. 1, CWI 84680, CWI 84694, CWI 84704, CWI 84686, CWI 14537, GEO-L1, WATDE0105, WATDE0944, WATDE0957, WATDE1005, WATDE1018, WATDE1017, WATDE0113, WATDE1010; 2, C33, WATDE1031, WATDE1032; 3, BW 50849, CWI 14244, CWI 28055, WATDE0026, WATDE0749, WATDE0047, WATDE0739, WATDE0999, WATDE1003, WATDE0993; 4, CWI 86929, CWI 86942, WATDE0975, WATDE0973, WATDE0974. The IBSpy variation values for the Watkins lines (WATDE) were extracted from Cheng *et al.* (2023).

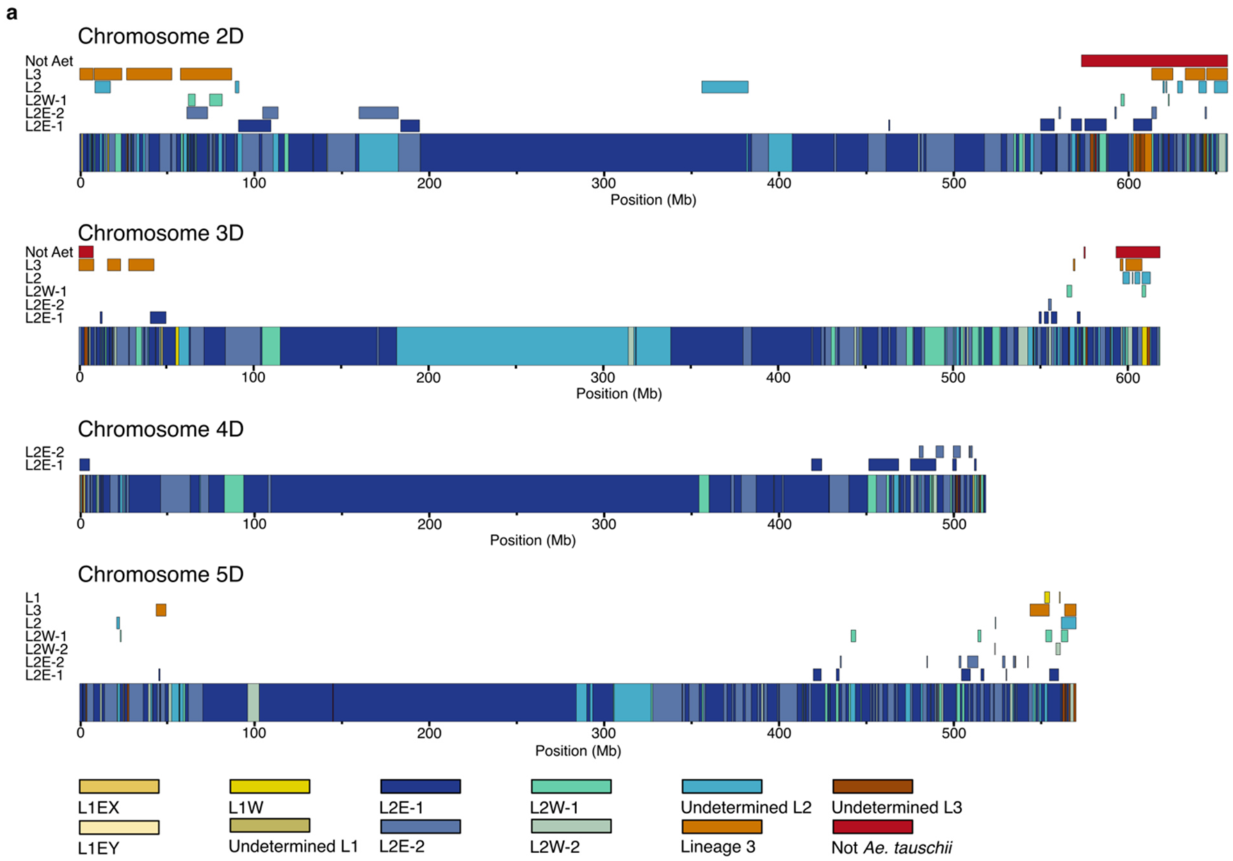


**Extended Data Fig. 7. Minimal number of hybridizations that gave rise to the extant bread wheat D genome**. Shown are graphical representations of Chinese Spring chromosomes 2D, 3D, 4D and 5D. The colored boxes in the chromosomes represent the haplotypes found in Chinese Spring. Colored rectangles above the chromosomes represent alternative haplotype blocks identified across 126 hexaploid wheat landraces (cumulative length of alternative haplotype blocks across all 126 landraces). Colors refer to the *Ae. tauschii* subpopulations following the legend. The maximum number of haplotype blocks is four.

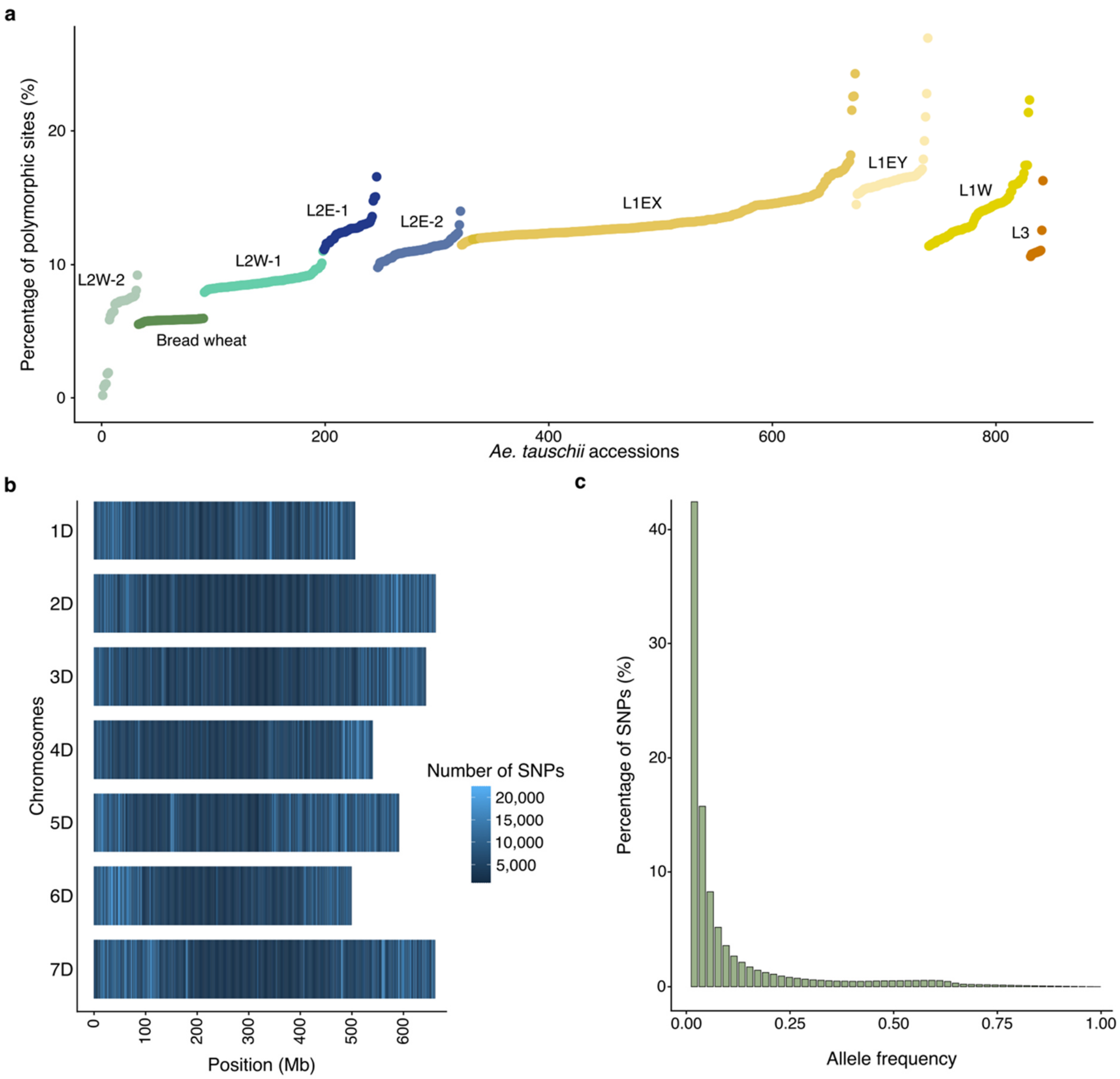


**Extended Data Fig. 8. SNP data statistics. a**, The percentage of polymorphic sites for each *Ae. tauschii* accession compared to the TA1675 (L2) reference accession. Each color represents an *Ae. tauschii* or bread wheat group. **b**, SNP density in windows of 1 Mb computed across the 7 chromosomes of TA1675. **c**, Allele frequency distribution.

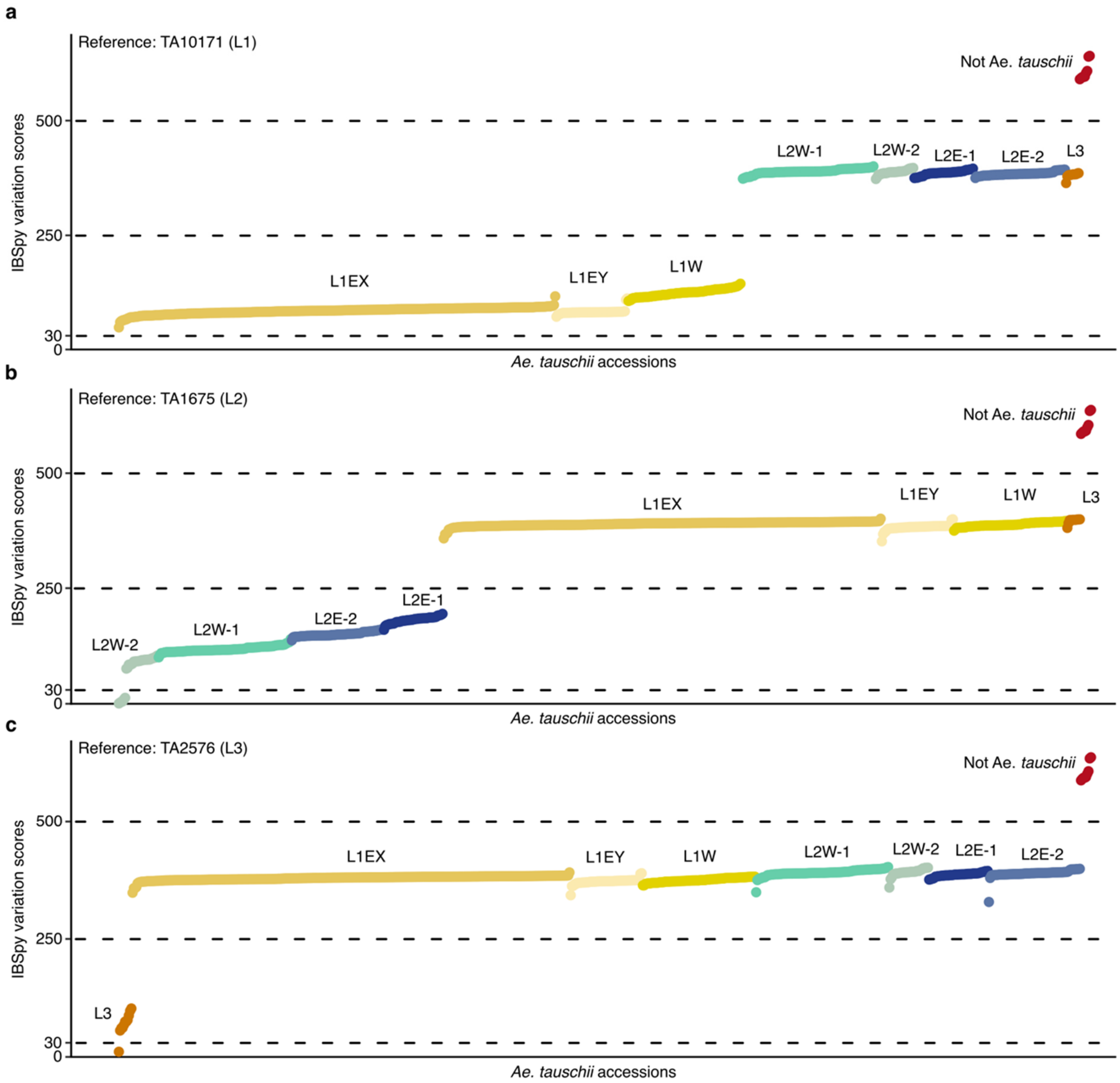


**Extended Data Fig. 9. IBSpy variation score distribution.** Shown are the average variation scores for each *Ae. tauschii* accession (represented as a dot) against TA10171 (L1) **(a)**, TA1675 (L2) **(b)**, and TA2576 (L3) **(c)** (Supplementary Table 28). Based on the distribution, we defined IBSpy values ≤30 as identical by state, values > 30 ≤ 250 as being the same *Ae. tauschii* lineage as the reference, values > 250 ≤ 500 as being a different *Ae. tauschii* lineage, and values > 500 as not being *Ae. tauschii*.

**Extended Data Table 1.** Assembly statistics for the three chromosome-scale *Aegilops tauschii* references and wheat landrace CWI 86942.

| Species | *Aegilops tauschii* | | | *Triticum aestivum* |
|---|---|---|---|---|
| Accession | **TA10171** | **TA1675** | **TA2576** | **CWI 86942** |
| Lineage | 1 | 2 | 3 | / |
| Sequencing coverage (fold) | 77 | 97 | 67 | 32 |
| Contig N50 (bp) | 53,375,588 | 221,041,983 | 116,906,157 | 44,458,675 |
| Assembly length (bp) | 4,151,983,908 | 4,159,914,615 | 4,245,074,256 | 14,571,138,882 |
| Pseudochromosomes | 7 | 7 | 7 | 21 |
| Length of pseudochromosomes (bp) | 4,106,536,600 | 4,106,562,375 | 4,124,033,060 | 14,470,226,287 |
| Number of unplaced scaffolds | 924 | 884 | 3,045 | 3,881 |
| Unplaced scaffold length (bp) | 45,447,308 | 53,352,240 | 121,041,196 | 100,912,595 |
| Unplaced scaffold N50 (bp) | 49,187 | 58,980 | 36,307 | 82,235 |
| Number of gapped regions | 131 | 32 | 79 | 878 |
| Number of HC gene models | 44,275 | 43,511 | 43,786 | 147,646* |
| QV (Phred score) | 40.93 | 48.06 | 40.3 | / |
| BUSCO score (%) | 98.02 | 98.33 | 98.14 | 99.1 |

*Number of lifted gene models
