## Supplementary notes for "Origin and evolution of the bread wheat D genome"

**Supplementary Note 1**

**Plant material**

The line/accession ID under which the line/accession was received, bulked, and sequenced is highlighted in bold text in Supplementary Table 1. Subsequent, previous, or parallel IDs assigned are also included based on indicated sources. Key germplasm bank IDs considered in this study are listed in columns C, D and E. Additional Germplasm bank/line/accession IDs, primarily from data obtained from genesys-pgr.org are listed in column F. Each ID may appear multiple times because collections are in part or completely duplicated between germplasm banks. Obvious errors such as circular references were identified in Genesys and removed; however, the full list of Additional IDs may contain unidentified historical errors and does not represent a full list of every ID ever assigned to every line. It is meant to be as comprehensive a record of the publicly available data as possible. It is up to the user to verify the provenance of any line they may use under the IDs listed within this table and its relevance to the IDs sequenced as part of this project.

Single seed descent lines obtained from particular accessions were assigned a unique plant ID with the identifier BW_XXXXX (column R). Each of these identities is traceable to an individual bulk of an accession ID as indicated in bold in columns C-E and to the seedbank IDs contained within the same. The exact plant ID deposited in the seedbank may be different to the plant ID used for sequencing but both are descended from the same accession.

**Supplementary Note 2**

***k*-mer matrix generation for the *Aegilops tauschii* diversity panel**

The process of generating *k*-mer frequency matrices in the context of GWAS analyses is computationally intensive both in terms of memory requirements and compute time. This computational bottleneck becomes more pronounced for larger panels with sizable genomes sequenced at high depth; a tendency that is recently enabled by the increasing affordability of high throughput sequencing. Typically, *k*-mer counts are obtained for each accession independently using a *k*-mer counting software such as Jellyfish. For this first phase, the counts can be computed in parallel, and saved in separate look-up hash tables (dictionaries), which can be subsequently queried as part of the matrix generation phase. Traditionally, merging the hash tables is performed in a brute-force fashion by iterating over the panel sequentially and incrementally inserting the *k*-mer entries and their counts for the corresponding accession, or assigning the count for the *k*-mer in question and the accession in process if the entry already exists. The number of hash table look-ups required for filling the matrix is equivalent to the size sum of all hash tables, and the size of the matrix grows with the panel size and the cardinality of the *k*-mer set. Nowadays, this can amount to hundreds of accessions with billions of *k*-mer entries, making this naive approach impractical to say the least. This computational cost is irreducible, but without parallel computation it becomes restricting. As part of this work, we developed a simple yet efficient scheme for parallelizing this workload, which we include here for the benefit of all.

Let us assume we have a panel of $N$ accessions $\{\mathcal{G}_1, \mathcal{G}_2, \ldots, \mathcal{G}_N\}$, and let $\mathcal{K}_i = \{\kappa, \kappa \in \mathcal{G}_i\}$ be the set of *k*-mers $\kappa$ pertaining to an accession $\mathcal{G}_i$. Also, let $c_j^i$ denote the frequency of *k*-mer $\kappa_j$ in accession $\mathcal{G}_i$:

Note that the *k*-mers do not assume any particular order and are indexed merely according to their arbitrary insertion order. Merging the first two accessions takes $|\mathcal{K}_1| + |\mathcal{K}_2|$ look-ups/fetches, and results in the $|\mathcal{K}_1 \cup K_2| \times 2$ partial matrix:

$$\begin{bmatrix} c_1^1 \\ c_2^1 \\ \vdots \\ c_{|\mathcal{K}_1 \cup \mathcal{K}_2|}^1 \end{bmatrix} \begin{bmatrix} c_1^2 \\ c_2^2 \\ \vdots \\ c_{|\mathcal{K}_1 \cup \mathcal{K}_2|}^2 \end{bmatrix}$$

Following an incremental process of incorporating one accession into the matrix at a time, the final matrix obtained through this process is:

$$\begin{bmatrix} c_1^1 & c_1^2 & \cdots & c_1^N \\ c_2^1 & c_2^2 & \cdots & \\ \vdots & \vdots & \ddots & \\ c_{\left|\bigcup_{i=1}^{N} \mathcal{K}_i\right|}^1 & & & c_{\left|\bigcup_{i=1}^{N} \mathcal{K}_i\right|}^N \end{bmatrix}$$

This matrix has a size:

$$\left|\bigcup_{i=1}^{N} \mathcal{K}_i\right| \times N$$

and the number of hash table look-ups required for its construction is:

$$\sum_{i=1}^{N} |\mathcal{K}_i|$$

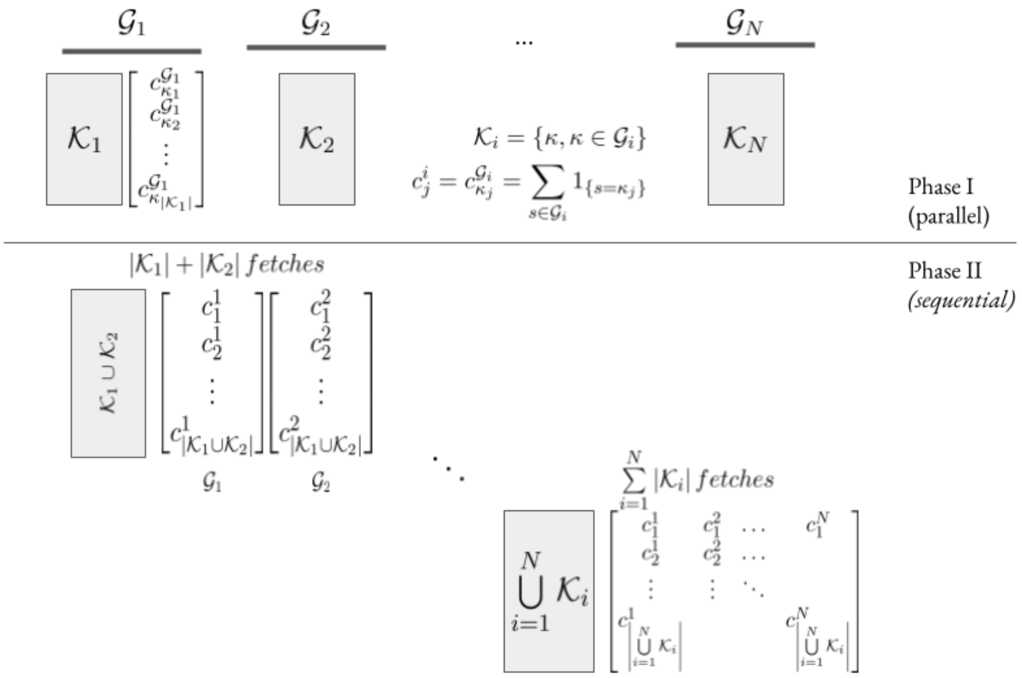


Brute-force computation of *k*-mer frequency matrix across a GWAS panel of N accessions

In order to parallelize this process, we introduce a hashing function $h$ which bins the binary encoding of a *k*-mer $\mathcal{E}(\kappa)$ to one of $m$ buckets:

$$h : \mathcal{K} \to \{1, 2, ..., m\}$$
$$\kappa \mapsto h(\kappa) = \mathcal{E}(\kappa) \mod m$$

The hash function introduces a partitioning $\mathcal{H}$ of the k-mer space:

$$\mathcal{H} : \mathcal{K} \to \{\mathcal{B}_1, \mathcal{B}_2, , \mathcal{B}_m\}$$
$$\kappa \mapsto \mathcal{H}(\kappa) = \mathcal{B}_{h(\kappa)}$$

This way an accession $\mathcal{G}_i$ is sharded into $m$ bins during the *k*-mer counting phase:

$$\mathcal{G}_i \xrightarrow{\mathcal{H}} \{\mathcal{B}^i_1, \mathcal{B}^i_2, ..., \mathcal{B}^i_m\}$$
$$\mathcal{B}^i_j = \{\kappa, \kappa \in \mathcal{K}_i \wedge h(\kappa) = j\}$$

Each bin $\mathcal{B}^i_j$ is a hash table indexing the counts of k-mers mapped to bin $j$ against accession $\mathcal{G}_i$. The obtained bins are mutually exclusive and their union represents a full k-mer count index of $\mathcal{G}_i$:

$$\bigcup_{j=1}^{m} \mathcal{B}^i_j = \mathcal{K}_i$$
$$\mathcal{B}^i_j \cap \mathcal{B}^i_{j'} = \phi, \forall j \neq j'$$

Consequently, it follows that:

$$\bigcup_{j=1}^{m} \mathcal{B}^i_j = \mathcal{K}_i$$
$$\mathcal{B}^i_j \cap \mathcal{B}^i_{j'} = \phi, \forall j \neq j'$$

More importantly, a *k*-mer is guaranteed to be mapped to the same bin index for all the accessions it belongs to, and the *k*-mer counts can now be looked-up from a pre-defined subset of bins, rendering the merger phase data parallel:

$$\kappa \in \mathcal{G}_i \cap \mathcal{G}_j \wedge \kappa \in \mathcal{B}^i_x \implies \kappa \in \mathcal{B}^j_x$$

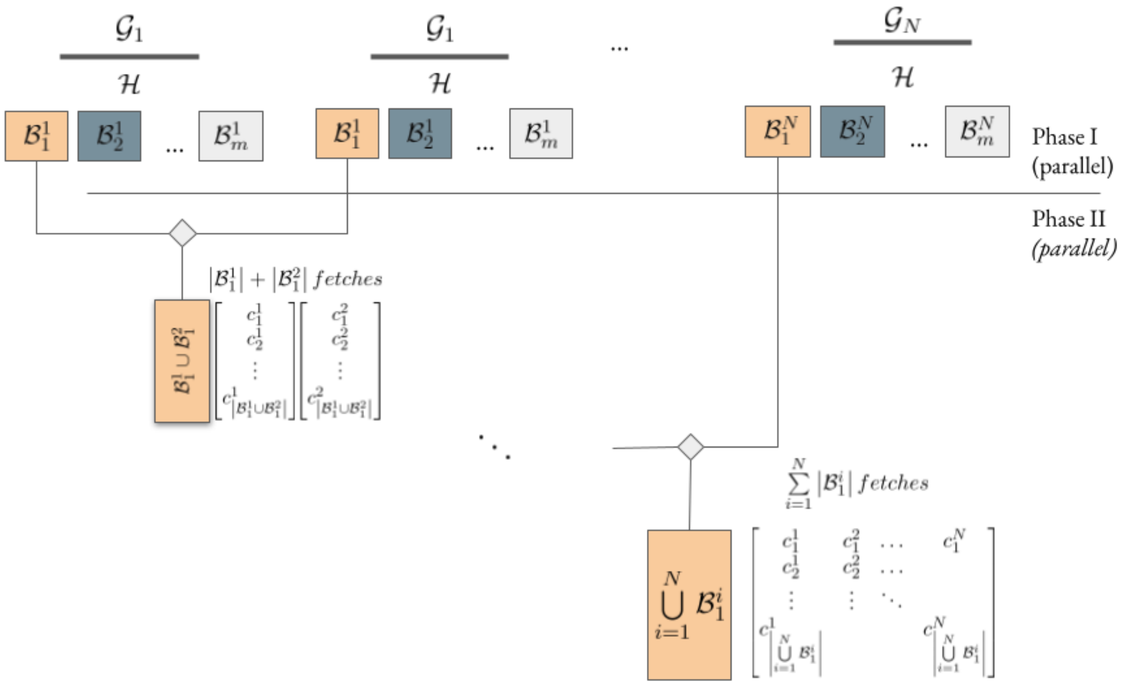


Parallelization of the k-mer frequency matrix generation through k-mer binning

For each bin index we assign a merger job which acts on the corresponding bins in a similar fashion as the brute-force approach described above. However, each job can run independently on much smaller data, requiring much less computation and memory. In particular, the number of hash table look-ups required per job is:

$$\sum_{i=1}^{N} \left|\mathcal{B}_j^i\right|$$

With a matrix size of:

$$\left|\bigcup_{i=1}^{N} \mathcal{B}_j^i\right| \times N$$

But how does that compare with the original approach?

Let $\overline{\mathcal{K}}$ denote the mean *k*-mer index size per accession:

$$\bar{\mathcal{K}} = \frac{1}{N}\sum_{i=1}^{N}|\mathcal{K}_i|$$

In GWAS, accessions usually pertain to closely related strains and are therefore similar in genome size. It follows from this that *k*-mer index size has low variance across accessions (regardless of whether the sequencing depth varies across the panel, which only affects the recorded frequencies):

$$|\mathcal{K}_i| \approx \overline{\mathcal{K}}, \forall i$$

Let $\overline{\mathcal{B}}^i$ denote the mean bin size obtained through $\mathcal{H}$ for accession $\mathcal{G}_i$:

$$\overline{\mathcal{B}}^i = \frac{1}{m}\sum_{j=1}^{m}|\mathcal{B}_j^i|$$

Because of the random nature of *k*-mer occurrence within a genome, which is for all practical considerations akin to a sampling from a uniform distribution of *k*-mer patterns, the hash function $h$ produces almost equally sized *k*-mer buckets for each accession, which, combined with the previous observations, gives:

$$\left|\mathcal{B}_j^i\right| \approx \bar{\mathcal{B}}^i = \frac{1}{m}|\mathcal{K}_i| \approx \frac{1}{m}\bar{\mathcal{K}}, \forall(i,j)$$

This is desirable, because it means the partitioning not only allows for the independent merger of same-index bins, but it also ensures that the computational load across the merger jobs has low variance. For example, in a fully parallel setting (where all jobs have immediate access to computational resources without queuing), the time it takes for the merger to finish depends on the worst performing job, and as we have demonstrated, this is not far from the average case, which is $m$ fold faster:

$$\sum_{i=1}^{N}\left|\mathcal{B}_j^i\right| = N\overline{\mathcal{B}}^i \approx \frac{N}{m}\overline{\mathcal{K}} = \frac{1}{m}\sum_{i=1}^{N}|\mathcal{K}_i|$$

Note that $m$ can theoretically be made arbitrarily large bar a few practical considerations, and we have had runs with $m > 1000$, enabling the computation of the matrix within mere minutes as opposed to more than a month using older code. The main consideration when deciding on the level of parallelism is whether the HPC resource can support the required IO throughput without introducing too much overhead. Note that the indexing phase would have to produce $mN$ files with $N$ parallel read/write tasks, while the merger phase would be merging $N$ files per job, with $m$ jobs running in parallel and performing read/write operations. With the advent of HPC storage and the increasing trend of incorporating cost-effective high performance solid-state storage solutions and supporting middleware to cater for the emerging data intensive applications, it is becoming more customary to leverage high throughput IO for achieving high data parallelism and reducing memory requirements per job.

We implemented this procedure as a C++ application (https://github.com/githubcbrc/KGWASMatrix) which comes with two binaries, one for producing a sharded *k*-mer count index, and a second for performing the merger step in order to produce the final matrix. For this study, we elected to perform our analysis based on the presence/absence binary matrix, however, the codes also allow for producing frequency matrices. The code is dockerised and the jobs can be run as singularity instances on HPC resources. The resource includes a brief documentation with details of how to run the codes, the various exposed parameters, and example template scripts for how to deploy this on an HPC cluster or a supercomputer.

**Supplementary Note 3**

***Aegilops tauschii* subpopulations contributing to the bread wheat D genome**

This approach was used to assess the origin of the bread wheat D genome across 17 hexaploid wheat chromosome-scale assemblies (Fig. 4b). Each chromosome-scale wheat assembly is divided into 50-kb windows and the origin of each window is determined based on identity-by-state to the *Ae. tauschii* subpopulations (at *K*=9). The pipeline can be found on GitHub (https://github.com/emilecg/wheat_evolution).

Input files. The files required as input are: (i) The output of the IBSpy runs. These are 17 tables (one for each hexaploid wheat line) showing the IBSpy values for each 50-kb window (row) against the 995 resequenced *Ae. tauschii* accessions (columns). (ii) A table specifying the corresponding subpopulation for each of the resequenced *Ae. tauschii* accessions based on 70% ancestry. (iii) Three lineage-specific tables (L1, L2, L3) specifying the corresponding subpopulation for the resequenced *Ae. tauschii* accessions based on 70% ancestry. The tables have four columns, the first containing the name of the accession, the second for the subpopulation at *K*=9, the third for the subpopulation at *K*=15 and the last for *K*=22. The input tables can be found here: (https://github.com/emilecg/wheat_evolution).

Determining identity-by-state. The script loops through each 50-kb window singularly to determine its origin in an independent way. The first step is to establish if there are *Ae. tauschii* accessions that show identity-by-state for any given 50-kb window. We used the following IBSpy variation values: ≤30 = identity-by-state, 31-250 = no identity-by-state, but lineage-specific (meaning that this 50-kb window can be assigned to a particular *Ae. tauschii* lineage, but not an accession), >500 = not *Ae. tauschii* (Extended Data Fig. 9, Supplementary Table 28).

If no *Ae. tauschii* accessions with identity-by-state were found for a given 50-kb window, the goal of the script is to determine which of the three lineages (L1, L2, L3) is the closest. This is done by calculating the proportions of L1, L2, and L3 accessions showing IBSpy variation values between 31 and 250. If no *Ae. tauschii* accession showed an IBSpy variation value below 250, the corresponding 50-kb window was considered to have an origin other than *Ae. tauschii* (introgression from other species).

For the 50-kb windows that showed identity-by-state to one or several *Ae. tauschii* accessions, the script determines the corresponding subpopulation to which these *Ae. tauschii* accessions belong. Similar to the approach above, this is in principle done by calculating proportions (i.e., number of *Ae. tauschii* accessions showing identity-by-state that belong to a particular subpopulation / total number of *Ae. tauschii* accessions defining this subpopulation). However, we realized that factors including the frequency of a given haplotype, the variable haplotype diversity across the chromosomes, and the varying numbers of accessions representing a subpopulation can increase the noise in our analysis. To reduce the noise, we introduced the following steps.

1) We observed that 50-kb windows for which many *Ae. tauschii* accessions showed IBSpy values ≤30 generally fall into regions with low genetic diversity (often pericentromeric regions that show lower genetic diversity between the subpopulations than telomeric regions). For such regions, the threshold of 30 might be too high. To increase the robustness, we ordered the *Ae. tauschii* accessions showing identity-by-state (lowest to highest IBSpy variation values for every 50-kb window). For every 50-kb window, we then determined the number of *Ae. tauschii* accessions ($y$) that are retained to assign a subpopulation. This number of accessions varies for every 50-kb

window and depends on the total number of *Ae. tauschii* accessions showing IBSpy variation values ≤30 ($x$).

$$y = \left(\sqrt{x} \times 2\right) + 10$$

The script only retains the selected number of accessions ($y$) with the lowest IBSpy variation values.

2) To account for the different numbers of *Ae. tauschii* accessions in each subpopulation, the proportion score was computed for each subpopulation in a non-linear way using a tangent function. The function has been adapted to give a high score when the majority of the *Ae. tauschii* accessions of a specific subpopulation are in the selected list. The population is assigned based on the highest score obtained with the following function where $x$ is the number of accessions for a specific subpopulation present in the selected list and $n$ is the total number of accessions in the subpopulation.

$$y = 2 \times (\frac{2\pi x}{2.3n} + \frac{\pi}{1.7})$$

We also assigned a robustness score for each 50-kb window, determined by the sum of the pairwise products of the combinations of all the scores. The maximum of the confidence is reached with a zero and the values are capped at 500, indicating high uncertainty of the subpopulation assignment. Most 50-kb windows have very low scores, indicating a high robustness of our analysis. Some pericentromeric regions can be harder to assign to a specific subpopulation, resulting in higher scores.

**Supplementary Note 4**

**Determining the minimal number of hybridizations that gave rise to the extant bread wheat D genome**

To determine the minimal number of hybridizations that gave rise to the extant bread wheat D genome, we assessed the maximum number of syntenic haplotype blocks across 126 resequenced bread wheat landraces (Supplementary Table 15). We decided to focus on landraces to exclude haplotype blocks that are the result of synthetic hexaploid wheat breeding.

First, IBSpy variant scores were calculated for all the 126 resequenced wheat landraces in 50-kb windows, using the Chinese Spring IWGSC RefSeq (v2.1) as reference. The variation values were visualized as a heatmap, which allowed us to visually determine alternative haplotype blocks (i.e., haplotype blocks that are diverged from Chinese Spring). Each landrace showing alternative haplotype blocks was retained for the following analysis. IBSpy provides a "divergence score", but does not indicate if alternative haplotype blocks found in two landraces at the same position have the same or different origins. To assess the origin (subpopulation) of alternative haplotype blocks, we produced IBSpy runs with the resequencing data of the 126 hexaploid wheat landraces against 46 chromosome-level *Ae. tauschii* assemblies. This allowed us to assign alternative haplotype blocks to one or multiple *Ae. tauschii* accessions and a subpopulation. Compared to the quantitative assessment described in Supplementary Note 3, the qualitative assessment done here was more straightforward and produced less noise (because we considered larger haplotype blocks).

Most alternative haplotype blocks showed complex patterns of recombinations, resulting in various lengths in different bread wheat landraces (see Fig. 3e for an example). For our analysis, we retained the largest cumulative block that could be assigned to a given *Ae. tauschii* subpopulation.
